# Longer mandible or nose? Co-evolution of feeding organs in early elephantiforms

**DOI:** 10.1101/2023.08.15.553347

**Authors:** Chunxiao Li, Tao Deng, Yang Wang, Fajun Sun, Burt Wolff, Qigao Jiangzuo, Jiao Ma, Luda Xing, Jiao Fu, Ji Zhang, Shi-Qi Wang

**Affiliations:** University of Chinese Academy of Sciences, Beijing 100049, China; Key Laboratory of Vertebrate Evolution and Human Origins of the Chinese Academy of Sciences, Institute of Vertebrate Paleontology and Paleoanthropology, Chinese Academy of Sciences, Beijing, China; Department of Earth, Ocean and Atmospheric Science, Florida State University, Tallahassee, FL 32306-4520, USA; Environmental Science & Technology, University of Maryland, College Park, MD 20742, USA; School of Civil and Hydraulic Engineering, Huazhong University of Science and Technology, Wuhan 430074, China; National Center of Technology Innovation for Digital Construction, Wuhan 430074, China

## Abstract

The long-trunked elephantids underwent a significant evolutionary stage characterized by an exceptionally elongated mandible. The initial elongation and subsequent regression of the long mandible, along with its co-evolution with the trunk, present an intriguing issue that remains incompletely understood. Through comparative functional and eco-morphological investigations, as well as feeding preference analysis, we reconstructed the feeding behavior of major groups of longirostrine elephantiforms. In the *Platybelodon* clade, the rapid evolutionary changes observed in the narial region, strongly correlated with mandible and tusk characteristics, suggest a crucial evolutionary transition where feeding function shifted from the mandible to the trunk, allowing proboscideans to expand their niches to more open regions. This functional shift further resulted in elephantids relying solely on their trunks for feeding. Our research provides insights into how unique environmental pressures shape the extreme evolution of organs, particularly in large mammals that developed various peculiar adaptations during the late Cenozoic global cooling trends.

## Introduction

Proboscideans are known for their exceptionally elongated and versatile trunks (*Shoshani, 1998*). However, unlike modern elephants, proboscideans underwent a prolonged evolutionary phase characterized by the presence of greatly elongated mandibular symphysis and mandibular tusks. This elongation can be traced back to the Late Oligocene species *Palaeomastodon* and *Phiomia*, which are among the earliest elephantiforms (*Andrews, 1906*), and continued through to the Late Miocene *Stegotetrabelodon*, a stem elephantid (*Shoshani, 1996*; *Tassy, 1996*). Extreme longirostriny, a feature observed in fossil and modern fishes, reptiles, and birds, was relatively rare among terrestrial mammals and its occurrence in large-bodied proboscideans is particularly intriguing. Particularly, during the Early and Middle Miocene (approximately 20–11 Ma), the morphology of mandibular symphysis and tusks exhibited remarkable diversity, with over 20 genera from six families (Deinotheriidae, Mammutidae, Stegodontidae, “Gomphotheriidae”, Amebelodontidae, and Choerolophodontidae) displaying variations (*Shoshani, 1996*; *Tassy, 1996*). Why did proboscideans have evolved such a long mandible of so diversified morphology? How did fossil proboscideans use their strange mandibular symphysis and tusks, and what was the role of trunk in their feeding behavior? Finally, what was the environmental background for the co-evolution of their mandible and trunks, and why did proboscideans finally lose their long mandible? These important issuers on proboscideans evolution and adaptation remain poorly understood. Addressing these significant aspects of proboscidean evolution and adaptation requires comprehensive investigations into the functional and eco-morphology of longirostrine proboscideans.

## Results

### Mandible and skull morphology of bunodont elephantiforms

During the Early and Middle Miocene, bunodont elephantiforms, as the ancestral group of living elephants, flourished (*Osborn, 1936*; *Gheerbrant and Tassy, 2009*; *Cantalapiedra et al., 2021*). Bunodont elephantiforms include Amebelodontidae, Choerolophodontidae and “Gomphotheriidae”; all possess a greatly elongated mandibular symphysis (*Gheerbrant and Tassy, 2009*; *Cantalapiedra et al., 2021*). Our comprehensive phylogenetic reconstructions contained the majority of longirostrine elephantiform taxa at the species level and strongly supported this taxonomic scheme (*Figure 1A, S1, Appendix S1, Data S1, Code S1–S3*). The three families were characterised by their distinctive mandible and mandibular tusk morphology (*Shoshani, 1996*; *Tassy, 1996*). The paraphyletic “Gomphotheriidae” have clubbed lower tusks (*Figures 1B, K, S2B*); their mandibular symphysis is relatively narrow. This morphology is rather unspecialised, and the extant elephants are derived from “Gomphotheriidae” (*Tassy, 1996*; *Cantalapiedra et al., 2021*). The mandibular symphysis of Amebelodontidae is generally shovel-like and the mandibular tusks are usually flattened and wide. *Platybelodon* is the most specialised genus within this family; it possesses extremely flattened and widened mandibular tusks with a sharp distal cutting edge (*Wang et al., 2013*) (*Figures 1C, I, J, S2A*). Choerolophodontidae is unique because it completely lacks mandibular tusks and it has a long trough-like mandibular symphysis (*Figures 1D, L, M, S2C–F*). A very deep slit is present on each side of the distal alveolar crest (distal mandibular trough edge) (*Figure 1L*), which is presumably for holding a keratinous cutting plate (*Figure 1M*), similar to the slits for holding large claws in felids and some burrowing mammals (e.g., anteaters). The anterior mandibular foramen of *Choerolophodon* is extremely large and tube-like (*Figure S2D*), which indicates a very developed mental nerve and eponymous artery (*Eales, 1926*), for the nutrition of tissues (i.e., keratins) grown from symphyseal dermis. The three lineages exhibit different evolutionary states of food acquisition organs (PC1 scores from mandible and tusk characters; see Supplementary Methods) (*Figure S3A, B*). Different food acquisition organs morphology strongly indicates different methods of food acquisition among the three gomphothere families.

**Figure 1.**
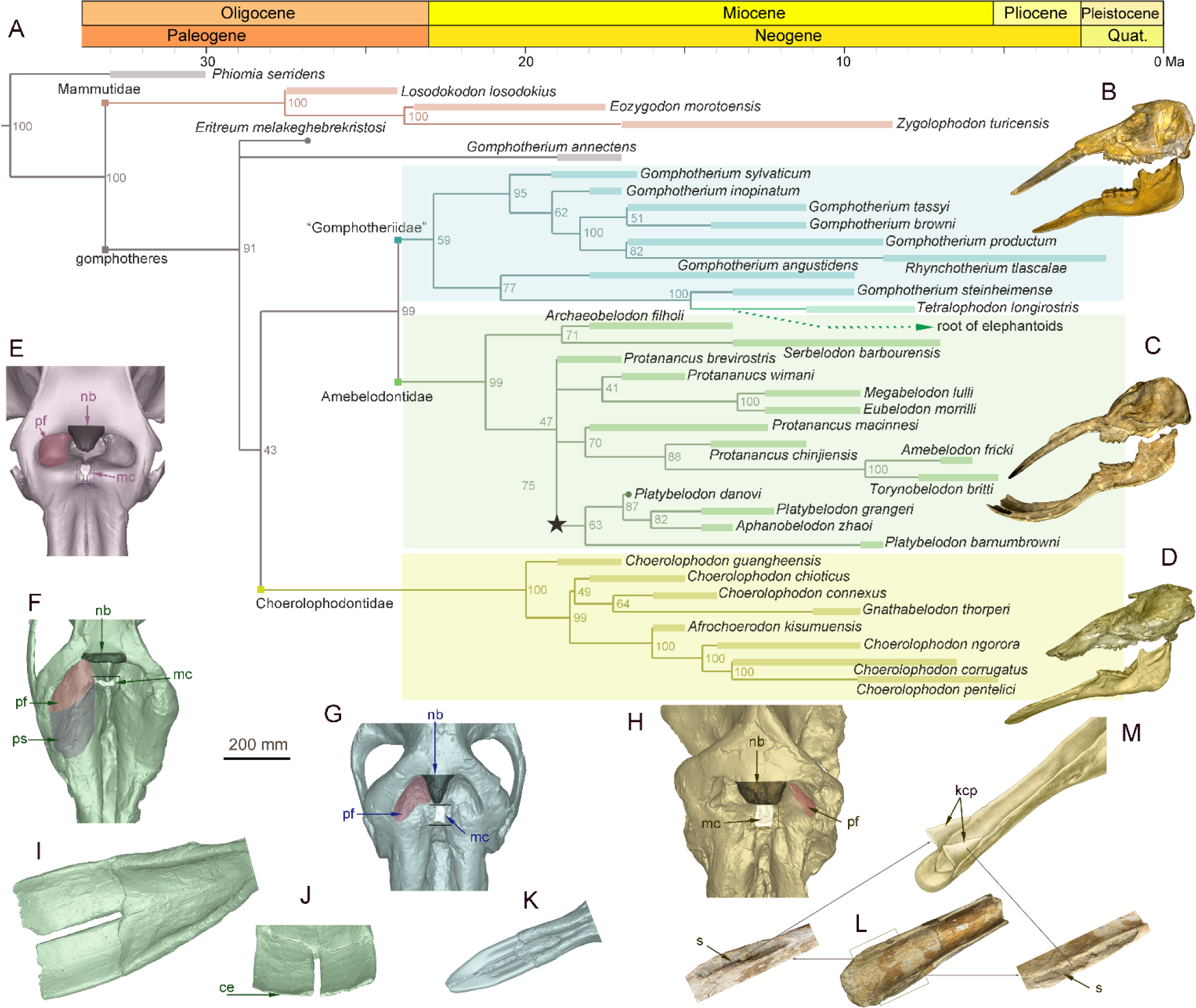
Morphology of the narial region and mandible of three gomphothere families compared with an extant elephant, and the elephantiformes phylogeny. (**A**) Phylogenetic reconstruction of major longirostrine elephantiforms at the species level based on the Bayesian tip-dating method. The node support (the number at each node) is the posterior probability, and the bars represent chronologic ranges of each taxon. (**B–D**) Representative cranium and mandible specimens of the three gomphothere families, including IVPP V22780, cranium, and IVPP V22781, mandible, of *Gomphotherium tassyi* [B], “Gomphotheriidae”, from Heijiagou Fauna, Tongxin region; HMV 0930, cranium and associated mandible of *Platybelodon grangeri* [C], from Zengjia Fauna, Linxia Basin; and IVPP V23457, cranium and associated mandible of *Choerolophodon chioticus* [D], from Middle Miaoerling Fauna, Linxia Basin. (**E–H**) Narial morphology of bunodont elephantiforms and elephantids in dorsal view, including IVPP OV733, *Elephas maximus* [E], a living elephantid; HMV 0930, *Platybelodon grangeri* [F]; IVPP V22780, *Gomphotherium tassyi* [G]; and IVPP V23457, *Choerolophodon chioticus* [H]. Mandibular morphology of bunodont elephantiforms. (**I–J**) Mandibular symphysis and tusks of HMV 0930, *Platybelodon grangeri*, in dorsal [I] and distal [J] views. (**K**) Mandibular symphysis and tusks of IVPP V22781, *Gomphotherium tassyi*, in dorsal view. (**L**) Mandibular symphysis of IVPP V25397, *Choerolophodon chioticus*, showing the deep slits at both sides of the distal alveolar crests in dorsal view. (**M**) Reconstruction of keratinous cutting plates in the slits, in dorsolateral view. Anatomic abbreviations: ce, cutting edge of the distal mandibular tusk in *Platybelodon*; kcp, reconstructed keratinous cutting plates in *Choerolophodon*; nb, nasal process of nasal bone; mc, slit or groove for mesethmoid cartilage insertion (white in color); pf, perinasal fossa; ps, prenasal slope in *Platybelodon*; s, slit for holding kcp in *Choerolophodon*.

### Evolutionary dynamics and ecological niches of three longirostrine bunodont elephantiforms families

The three longirostrine bunodont elephantiforms families were widely distributed in the Early–Middle Miocene in northern China, ranging from ∼19 to 11.5 Ma (*Figure 2, S4*) (most of the Shanwangian and Tunggurian stages of the Chinese Land Mammal Age) (*Deng et al., 2018*). However, the relative abundances of these three families were very different and varied through time. We investigated fossil bunodont elephantiforms from four regions: Linxia Basin (LX), Tongxin region (TX), Junggar Basin (JG), and Tunggur region (TG) (*Figures 2, S4, Data S2*). We focused on different fossil assemblages in different ages. We evaluated the amounts of fossils of each elephantiform taxon from major fossil assemblages in the above regions from three museums (IVPP, HPM and AMNH, for full names, please see the Material and Methods) and calculated their proportions among all bunodont elephantiforms fossils (*Figure 2A*). During ∼19–17 Ma, the quantities of both *Choerolophodon* (Choerolophodontidae) and *Protanancus* (Amebelodontidae) were larger than those of *Gomphotherium* (“Gomphotheriidae”) despite the high diversity of *Gomphotherium* in species level. During ∼17–15 Ma (the Mid–Miocene Climate Optimum [MMCO]), the relative fossil abundances of the three gomphothere families were similar. However, within Amebelodontidae, *Platybelodon* appeared and rapidly replaced the primitive *Protanancus*. After ∼14.5 Ma (i.e., the beginning of the Mid–Miocene Climate Transition [MMCT]) (*Westerhold et al., 2020*), *Choerolophodon* suddenly experienced regional extinction, and *Gomphotherium* also gradually declined. Within Amebelodontidae, *Aphanobelodon*, which is morphologically similar to *Platybelodon*, only occurred in one fossil assemblage, and the *Platybelodon* population greatly expanded. After ∼13 Ma, *Gomphotherium* were regionally extinct, and *Platybelodon* became the only genus among bunodont elephantiforms.

**Figure 2.**
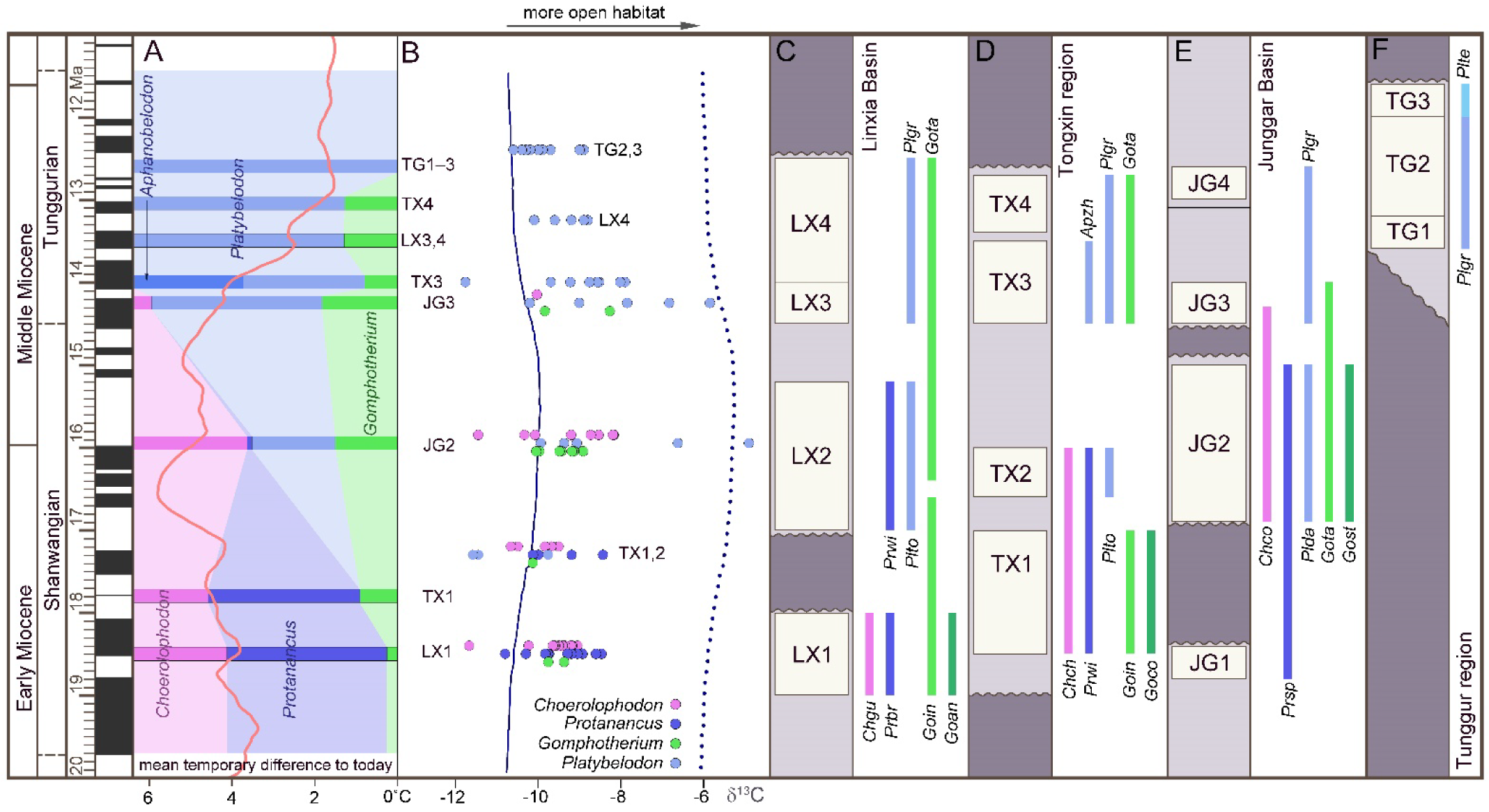
Relative abundance, tooth enamel δ^13^C, and stratigraphic ranges of the three gomphothere families in the Shanwangian and Tunggurian stages (∼20–11.5 Ma) of northern China. (**A**) Relative abundance of the three gomphothere families, including Choerolophodontidae, only represented by *Choerolophodon* (pink); Amebelodontidae, represented by *Protanancus*, *Aphanobelodon* (dark blue), and *Platybelodon* (light blue); and “Gomphotheriidae”, only represented by *Gomphotherium* (green). Horizontal bars indicate the average ages of the fossil assemblages, which are shown in C–F. The ages were determined by paleomagnetism (*Table S1*). The red curve shows the global reference benthic foraminifer oxygen isotope curve, which represents the global temperature [after (*Westerhold et al., 2020*)]. (**B**) Tooth enamel stable carbon isotopic compositions of various gomphothere taxa. Each circle represents the bulk enamel δ^13^C values of a single tooth. The data of LX4 and TG2,3 are from previous publications (*Wang and Deng, 2005*; *Zhang et al., 2009*). Solid and dashed lines represent the mean and maximum enamel δ^13^C values for C_3_ diets that have been corrected for Miocene atmospheric CO_2_ δ^13^C [after (*Tipple et al, 2010*)]. (**C–F**) Synthetic stratigraphic columns of typical fossil bearing regions of northern China during ∼19–11.5 Ma, which incorporated different fossil assemblages with different ages, from the Linxia Basin [C], Tongxin region [D], Junggar Basin [E], and Tunggur region [F]. Vertical bars represent the temporal ranges of different gomphothere taxa. For fossil assemblage abbreviations, please see Table S1. Abbreviations for gomphothere taxa: Apzh, *Aphanobelodon zhaoi*; Chch, *Choerolophodon chioticus*; Chco, *Choerolophodon connexus*; Chgu, *Choerolophodon guangheensis*; Goan, *Gomphotherium* cf. *angustidens*; Goco, *Gomphotherium cooperi*; Goin, *Gomphotherium inopinatum*; Gost, *Gomphotherium steinheimense*; Gota, *Gomphotherium tassyi*; Plda, *Platybelodon dangheensis*; Plgr, *Platybelodon grangeri*; Plte, *Platybelodon tetralophus*; Plto, *Platybelodon tongxinensis*; Prbr, *Protanancus brevirostris*; Prsp, *Protanancus* sp.; Prwi, *Protanancus wimani*.

We analyzed stable isotope values of the tooth enamel for each gomphothere taxon in different fossil assemblages (*Data S3*). Overall, the δ^13^C values indicate a relatively open environment that consisted of a diverse range of habitats (including grasslands, wooded grasslands, and forests) dominated by C_3_ plants in northern China from ∼19 to 11.5 Ma (*Figures 2B, S4*). In LX1 (*Figure 2B, Data S2*), at approximately ∼18.5 Ma (see *Table S1*), both δ^13^C and δ^13^O values of *Choerolophodon* and *Protanancus* are similar and have a wide range of overlap, and those of *Gomphotherium* are in the middle of the range.

This indicates that the niches of these three groups overlapped without obvious differentiation. In TX1,2 (∼17.3 Ma), during which *Platybelodon* first appeared, the δ^13^C values of *Platybelodon* are lower than those of *Protanancus*. The δ^13^C values of *Choerolophodon* and *Gomphotherium*, however, are all within the δ^13^C range of Amebelodontidae. In JG2 (∼16 Ma), the δ^13^C value of *Platybelodon* shows a distinct positive shift, which indicates expansion into more open habitats (*Figure 2B*). However, *Choerolophodon* may have persisted in a relatively closed environment (*Figure 2B*). Moreover, the ecological niche of *Gomphotherium* appeared to be in between those of *Platybelodon* and *Choerolophodon*, and they potentially lived in the boundary area of open and closed habitats (*Figure 2B*). This isotopic niche pattern is also observed in JG3 (∼14.3 Ma), with the rise of *Platybelodon* and decline of *Choerolophodon* and *Gomphotherium*. In TX3, LX4, and TG2,3 (after ∼14 Ma), only *Platybelodon* were examined (as *Gomphotherium* specimens are too rare to sample); its ecological niche was similar to that previously occupied by *Choerolophodon* after the latter went extinct.

### Specialized feeding behaviors of three gomphothere families

To reconstruct the feeding behaviors of the three gomphothere families, we performed Finite Element (FE) analyses on three models as representatives of each family: *Choerolophodon*, *Gomphotherium*, and *Platybelodon* (for model settings see *Figures S5–S8, Tables S2–S4, 3D models S1–S6*). We conducted two kinds of tests: the distal forces test (*Code S4*) and the twig-cutting test (*Code S5*). In the distal forces test, after full muscle forces were exerted, a 5000 N vertical force is loaded on the distal end of each mandible (*Figures 3A–C, Movies S1–S3*). The mandible strain energy curve (MSEC) of *Platybelodon* suddenly reaches a very high value, while the MSEC increases of *Gomphotherium* and *Choerolophodon* are far less than that of *Platybelodon* (*Figure 3D*). Then, keeping the magnitude of the external force unchanged, with change of the external force from vertical to horizontal direction, the MSEC of *Platybelodon* decreases even lower than that of *Gomphotherium* and was similar to that of *Choerolophodon*. This result indicates that the mechanical performance of the *Platybelodon* mandible is disadvantageous under distal vertical external forces, but greatly improved under horizontal external forces (*Figure 3A–C, Movies S1–S3*).

**Figure 3.**
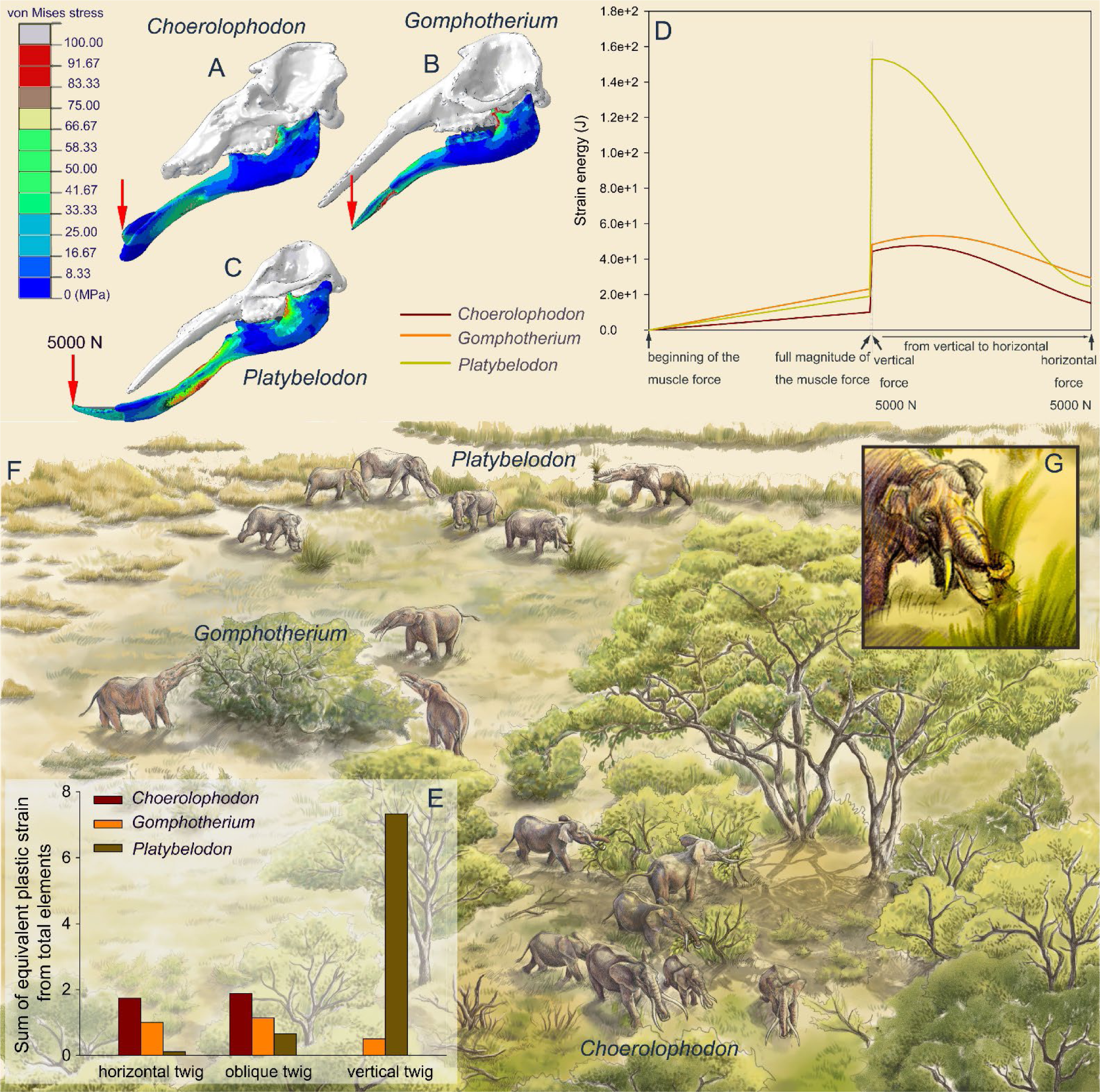
Finite element analyses of feeding behaviors among three longirostrine gomphothere families and reconstruction of their feeding ecology. (**A–C**) von Mises stress color maps of *Choerolophodon*, *Gomphotherium*, and *Platybelodon* models, with the full muscle forces exerted, and an additional 5000 N external vertical force applied on the distal end of the mandibular symphysis. (**D)** Strain energy curves of the three mandibles under the following three steps: 1, muscle forces linearly exerted; 2, a 5000 N external vertical force suddenly applied on the distal end; and 3, the 5000 N external force gradually changed from vertical to horizontal. (**E)** Sum of equivalent plastic strain from total elements (SEPS) of twigs cut by mandible models in three different directions (i.e., twig horizontal, 45° oblique, and vertical). Larger SEPS values indicate higher efficiency of twig cutting. (**F)** Scenery reconstruction of feeding behaviors of the three longirostrine gomphothere families (by X. Guo), represented by *Choerolophodon* (Choerolophodontidae), feeding in a closed forest, *Gomphotherium* (“Gomphotheriidae”), feeding at the margin between the closed forest and open grassland, and *Platybelodon* (Amebelodontidae), feeding on open grassland. (**G)** Detailed 3D reconstruction of *Platybelodon* feeding by grasping the grass-blades using their flexible trunk and cutting the grass blades using the distal edge of their mandibular tusks.

In twig-cutting tests, a cylindrical twig model of orthotropic elastoplasticity was posed in three directions to the distal end of the mandibular tusks of *Platybelodon* and *Gomphotherium*, and to the keratinous cutting plate of *Choerolophodon*. The sum of the equivalent plastic strain (SEPS) from total twig elements was calculated (equivalent plastic strain represents the irreversible deformation of an element, and the sum from all twig elements can reflect the cutting effects) (*Figure 3E*), and the cutting movies are provided (*Movies S4–S19*). When the twig was placed horizontally, the SEPS of the *Choerolophodon* model is the largest, which means that *Choerolophodon* has the highest twig-cutting efficiency, followed by *Gomphotherium*; while *Platybelodon* exhibits the much lower efficiency in cutting horizontal twigs than the other two models. When the twig was placed obliquely (45° orientation), the *Platybelodon* model still shows the smallest SEPS value, although it was nearly one order of magnitude higher than that of the horizontal twig. Finally, when the twig itself was in a vertical direction, the growth direction of the keratinous cutting plate determines that *Choerolophodon* cannot cut in this condition. The SEPS of the *Gomphotherium* model also decreases and shows lower cutting efficiency. In contrast, the SEPS of the *Platybelodon* model increases another order of magnitude, which is substantially larger than that of any other taxa in any cutting state. These data strongly indicate that *Platybelodon* mandible is specialized for cutting vertically growing plants. However, the *Choerolophodon* mandible is specialized for cutting horizontally or obliquely growing plants, this explains the absence of mandibular tusks, but they are likely not able to feed on vertically growing plants. The cutting effect of the *Gomphotherium* mandible is relatively even for all directions.

### Co-evolution of narial morphology and characters of horizontal cutting among gomphothere families

The three lineages with different mandibular morphology also exhibited different stages of trunk evolution, which can be inferred from the narial region morphology (*Tassy, 1994*). Among the three groups, *Gomphotherium* has similar narial morphology to living elephantids (*Figures 1E, G, S9D, E*), with comparable size and morphology of the nasal bone process, insertion for mesethmoid cartilage, and perinasal fossae. Alternatively, the narial region of *Choerolophodon* shows a relatively primitive evolutionary stage (*Tassy, 1994*) (*Figure 1H, S9A–C*). It has a very wide and large nasal bone process for an elephantiform, a fairly wide and long groove for mesethmoid cartilage insertion, and a pair of somewhat incipient (or even absent) perinasal fossae. Amebelodontids usually possess common narial morphology similar to that of *Gomphotherium* (*Tassy, 1994*). However, *Platybelodon* has noteworthy differences in narial morphology (*Wang and Li, 2022*) (*Figure 1F, S9G–L*). Its nasal aperture is greatly enlarged, which results in a very broad area for attaching *maxillo-labialis* (*Boas and Paulli, 1908*; *Eales, 1926*), the “core” muscle of the proboscis. The nasal bone process is very short and stout, with a slight dorsal bulge. The slit for mesethmoid cartilage insertion is very tiny. Beside the well-developed perinasal fossa, a vast inclined region is positioned rostral to the perinasal fossa, this is hereafter referred to as the prenasal slope (*Wang and Li, 2022*), and it potentially provides additional attachment for the *nasialis*.

The narial morphology of *Platybelodon* (and the closely related genus *Aphanobelodon* (*Wang et al., 2017*) (*Figure S9F*)), is unique in elephantiforms, and shows even more derived characters than living elephants. On the phylogenetic tree (*Figure 4A*), the *Platybelodon* lineage (the dark purple star in *Figure 4*) displays the most derived narial morphological combination (PC1 scores from characters in relation to the narial region), while Choerolophodontidae (the dark purple circle in *Figure 4*) show the least specialized narial morphology, which is close to the stem taxon *Phiomia*. Interestedly, the evolutionary level of the character-combine in relation to horizontal cutting (in terms of PC2) is highly correlated with that of narial region (*Figure 4B*). In the *Platybelodon* clade, this character-combine also shows high evolutionary level; while in the Choerolophodontidae clade, the evolutionary level of horizontal cutting is rather low, comparable with that of narial region. The result strongly suggests a highly co-evolution between narial region and horizontal cutting behavior in the trilophodont longirostrine bunodont elephantiforms.

**Figure 4.**
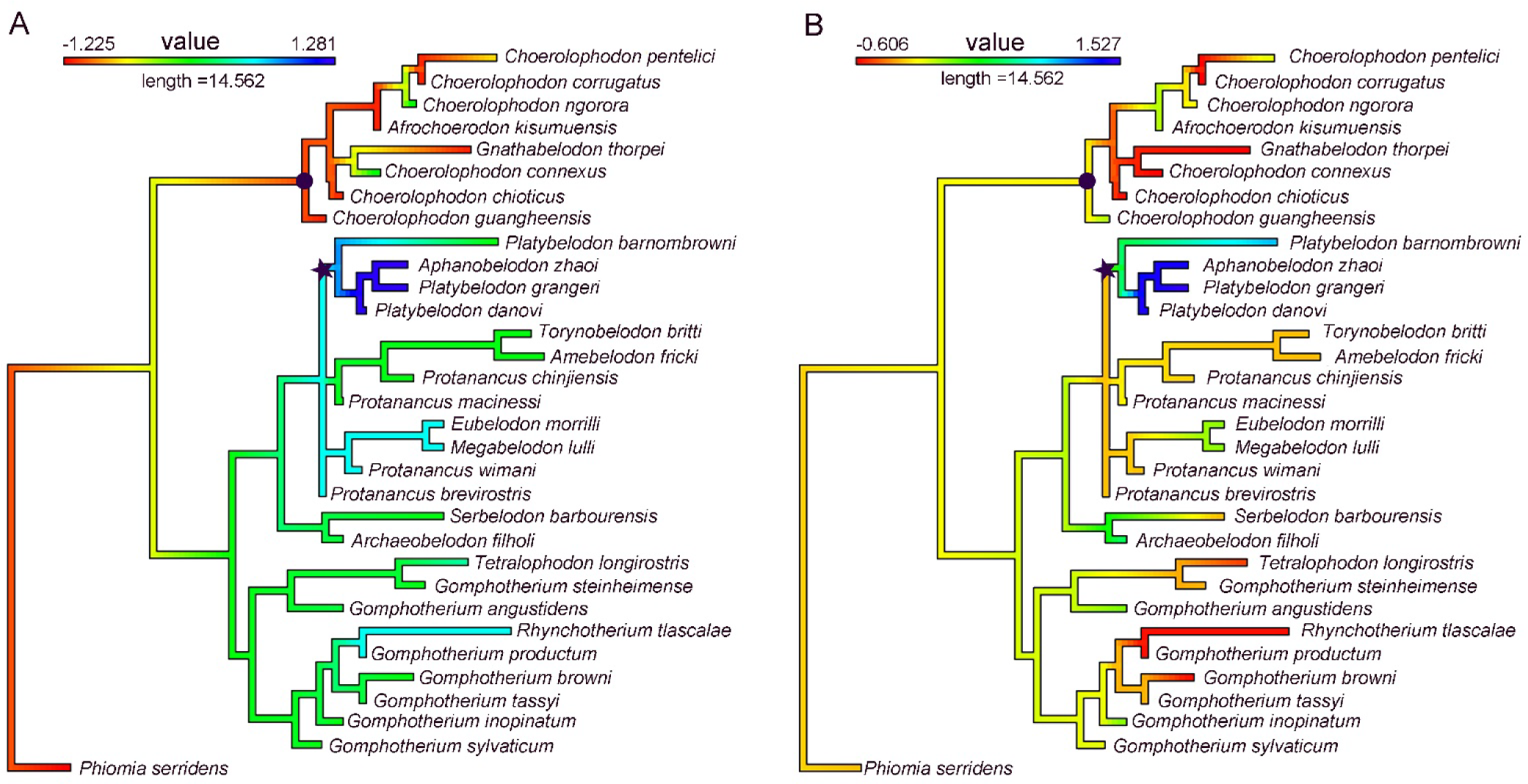
Evolutionary levels of narial region (**A**) and of characters in relation to horizontal cutting (**B**). Value in A was PC1 of characters 54–57 (*see Supplementary Appendix S1, Figure S3C, and Data S1*); and that in B was PC2 of characters 5, 9, 11, 72, and 77 (*see Figure S3D*). Note that the clade of *Platybelodon* (marked by an asterisk) shows high evolutionary levels and that of Choerolophodontidae (marked by a circle) shows low evolutionary levels in both character-combines, strongly suggesting the co-evolution of narial morphology and horizontal cutting behavior.

## Discussion

In several fossil and living terrestrial mammalian groups, including living *Tapirus* (*Moyano and Giannini, 2017*) and extinct *Proboscidipparion* (*Ma et al., 2023*), *Astrapotherium* (*Kramarz et al., 2019*), and *Macrauchenia* (*Blanco et al., 2021*), elongated noses have evolved as food procuring organs. However, none of these groups possess a long and dexterous trunk like elephants, as they have not lost their mandibular incisors. Only living elephants are capable of solely accomplishing food procurement with their trunks. Coincidentally, longirostrine proboscideans, characterized by their extremely elongated mandibular symphysis and tusks, are the only group where these highly developed organs may have co-evolve (*Tassy, 1996*). It is evident that the great elongation of the nose is well-matched with the extreme elongation of the mandibular symphysis. However, different lineages exhibit different evolutionary strategies strongly influenced by their ecological adaptations.

Our focus is on the ancestral elephantids, longirostrine bunodont elephantiforms, which include three families (Amebelodontidae, Choerolophodontidae, and “Gomphotheriidae”). These groups flourished during the MMCO, a period of global warmth from 17 to 15 Ma (*Zachos et al., 2001; Westerhold et al., 2020*). With the climatic shift during the MMCT (after ∼14.5 Ma), *Choerolophodon* sharply declined and experienced regional extinction in northern China. *Gomphotherium* also declined but persisted until around 13 Ma, while *Platybelodon* became predominant until the end of the Middle Miocene (*Figure 2, S4*). Worldwide, *Choerolophodon* and *Gomphotherium* continued to flourish during the MMCT and survived to the early Tortonian in other regions (*Lambert and Shoshani, 1998; Göhlich, 1999; Sanders et al., 2010*). However, *Platybelodon* was mostly restricted to Central Asia, especially northern China, with only a few records found in other regions (*Wang et al., 2013; Wang and Li, 2022*) (*Figure S10*). These uneven distributions were likely influenced by climatic and ecological factors, such as the relatively drier climate and more open ecosystems in northern China, and Central Asia, strongly affected by the elevation of the Tibetan Plateau (*An, et al., 2001*; *Wu et al., 2022)*. Extensive studies, using evidence from various disciplines including geochemical and geomagnetic proxies, and fossil records, has been conducted to explore this issue (*Jiang et al., 2008*; *Guo et al., 2002; Miao et al., 2012*; *Tang and Ding, 2013*; *Wang et al., 2022*).

As discussed, the three groups can be distinguished by different mandibular symphysis and tusk morphologies (*Gheerbrant and Tassy, 2009*), which indicate different feeding behaviors. Their distinct narial regions further reflect different evolutionary stages of their trunks (*Tassy, 1994*). *Choerolophodon* possesses a highly specialized mandibular symphysis for horizontally cutting growing plants and was confined to relatively close habitats. The low evolutionary level of the narial region suggests a relatively primitive or clumsy trunk in *Choerolophodon*. The feeding strategy of *Gomphotherium* was unspecialized and flexible, relying on the coordination of the enamel band of an upper tusk and the corresponding lower tusk. Previous research also suggested that *Gomphotherium steinheimense*, the sister taxon of elephantoids (*Figures 1A, S1*) from the Halamagai Fauna, fed on grasses (*Wu et al., 2018*). While FE analysis could not provide a clear suggestion for the trunk function of *Gomphotherium*, the narial evolutionary stage, which is close to living elephants, suggests a relatively flexible trunk in *Gomphotherium*, as it is phylogenetically closer to living elephants than the other two groups.

Enamel isotope results also support the idea that *Platybelodon* expanded its living habitats to more open environments, such as grasslands, more than any other bunodont elephantiforms, likely due to its distinct feeding strategy. Our FE analyses strongly indicate that the *Platybelodon* mandible is specifically suited for cutting vertically growing plants. In open environments, there are vertically growing plants, such as soft-stemmed herbs. *Platybelodon* did not survive to the Late Miocene in northern China, possibly due to a mass extinction caused by the global climatic shift known as the Tortonian Thermal Maximum. However, from another perspective, if the *Platybelodon* trunk functioned similarly to that of living elephants, such as pulling out herbs from the earth (*McKay, 1973*), the greatly enlarged mandibular symphysis and tusks would have become redundant.

The reduction of the mandibular symphysis and tusks in the Late Miocene occurred in every lineage of elephantiform, coinciding with the large-scale expansion of C4 grasses in the middle and low latitudes (*Cerling et al., 1997*). This may reflect the functional evolution of trunk grasping and manipulation in all elephantiforms lineages. From around 8 Ma to 5 Ma in Africa, some derived members of “Gomphotheriidae” (e.g., *Anancus*) and stem taxa of Elephantidae (e.g., *Stegotetrabelodon*, *Primelephas*) showed a strong inclination towards grazing on C4-grasses, even though their cheek tooth morphology remained relatively primitive (*Lister, 2013*). Thus, open environments might be a key factor in both trunk development and the evolution of modern elephants.

As we have discussed, mandibular elongation was a prerequisite for the extremely long trunk of proboscideans, and open-land grazing further promoted the evolution of trunks with complex manipulative functions. This may explain why tapirs never developed a trunk as dexterous as that of elephants, as tapirs never shifted their adaptation zones to open lands. Living in dense forests, foods were easily accessible and procured through the mouth. Furthermore, the living mammals of the Late Cenozoic, in various open areas, have undergone specific organ evolution, such as the elongated necks of giraffes (*Wang et al., 2022*), the extravagate saber-tooth evolution in carnivores (*Jiangzuo et al., 2023*), and the development of various strange cranial appendages in different ruminants (*Janis, 1982*). It is possible that the highly evolved trunks of elephants evolved somewhat accidentally, under the pressure of ecological changes from closed to open environments.

### Conclusion

In this study, we have examined the functional and eco-morphology, as well as the feeding behaviors, of longirostrine bunodont elephantiforms. Our findings demonstrate that multiple eco-adaptations have contributed to the diverse mandibular morphology observed in proboscideans, while open-land grazing has driven the development of trunk coiling and grasping functions and ultimately led to the loss of the long mandible. Specifically, the longirostrine elephantiform, *Platybelodon*, represents the first known proboscidean to have evolved both grazing behavior and trunk coiling and grasping functions. We have arrived at this conclusion through three lines of evidence, including the palaeoecological reconstruction based on tooth enamel stable isotope data, the reconstruction of feeding behaviors through finite element analyses, and the examination of mandibular and narial region morphology correlated with characteristics associated with horizontal cutting behavior. The coiling and grasping ability of the trunk in *Platybelodon* evolved within the ecological context of Central Asia, which experienced regional drying and the expansion of open ecosystems following the MMCT (*Miao et al., 2012*; *Tang and Ding, 2013*). As a result, *Platybelodon* outcompeted other longirostrine bunodont elephantiforms and flourished in the open environment of northern China until the end of the MMCT. This scenario sheds light on how proboscideans overcame an evolutionary bottleneck. Initially, the elongation of mandibular symphysis and tusks served as the primary feeding organs, with the trunk being used as an auxiliary tool. However, through some necessary modifications such as tactile specialization and water intake adaptations (*Shoshani, 1998; Purkart et al. 2022*), the feeding function gradually shifted entirely to the trunk, which offered advantages in terms of flexibility and lighter weight of the feeding organs. Consequently, elephantiforms rapidly reduced the length of their mandibular symphysis and tusks (*Shoshani and Tassy, 2005*; *Gheerbrant and Tassy, 2009*). Similar stories of open-land adaptation and the acquisition of iconic characteristics have been observed in various mega-mammalian lineages, highlighting the crucial role of open-land adaptation for successful survival in modern ecosystems (*Janis, 1982*).

## Methods

All examined crania and mandibles of the three longirostrine gomphothere families from the Early and Middle Miocene of northern China were housed in three museums: the Institute of Vertebrate Paleontology and Paleoanthropology (IVPP), American Museum of Natural History (AMNH), and Hezheng Paleozoological Museum (HPM, HMV is the specimen prefix). For FE analyses, surface 3D digital models of representative specimens were generated using a handheld Artec Spider 3-dimensional scanner, and the 3D models were reconstructed using Artec Studio 14 Professional. The geometrical models were further loaded into Abaqus CAE (V 6.14) for FE analyses. Phylogenetic reconstructions of longirostrine elephantiforms were performed using Bayesian total-evidence dating and maximum parsimonious analyses, which were done by MrBayes 3.2.7 and TNT 1.1, respectively. Principal component analyses were conducted using Past 4.04. Enamel isotopes analyses were conducted at the National High Magnetic Field Laboratory, USA. For detailed information, please see Supplementary Information.

## Supporting information

Models

## Acknowledgements

We thank the financial support received during the field work of the Second Tibetan Plateau Scientific Expedition. We appreciate Jie Ye, Xiaoxiao Zhang for his great guidance and suggestions on taxonomy and stratigraphy; Chi Zhang for assistance in phylogeny; Sijian Xu for assistance in fossil preparation; Jin Meng of AMNH, Rong Yang and Shanqin Chen of HPM for access to specimens in these two museums for comparison and statistics; Xiaocong Guo and Yu Wang for reconstruction. We thank Mallory Eckstut, PhD, from Liwen Bianji (Edanz) (www.liwenbianji.cn) for editing the English text of a draft of this manuscript. This work was supported by the National Key Research and Development Program of China (No.2023YFF0804501), National Natural Science Foundation of China (52178141, 41625005), National Science Foundation Cooperative Agreement No. DMR-1157490, the State of Florida.

## Author Contributions

S W, C L, T D, Y W, J Z designed the project; C L, S W, J F, performed 3D reconstruction; C L, S W, Q J, L X performed cladistic analyses; S W, J Z, C L, performed FEA; C L, Y W, F D, B W, J M determined the isotopes; C L, S W wrote the initial version of the paper; The all authors discussed the results and wrote the manuscript.

The authors declare that they have no competing interests.

## Additional Information

Supplementary Information is available for this paper, including material and methods, additional results and discussion, as well as the source data (Figures S1–S9, Tables S1–3, Appendix S1, Data S1–S4, Movies S1–S19, Codes S1–S5, and 3D Models S1–S6).

Reprints and permission information is available at www.nature.com/reprints

## Data availability statement

All the raw data for statistics and analyses are available in the Supplementary Information.

## Code availability statement

All the codes for analyses are available in the Supplementary Information.

## Notes

### Competing Interest Statement

The authors have declared no competing interest.

### Summary of Updates

1. We have replaced gomphotheres with "bunodont elephantiforms" throughout the entire manuscript as per the reviewer's suggestion. 2. Certain references have been corrected. 3. We modified the sequence of figures in the supplementary file, and Figure S3 underwent revisions. 4. Several minor adjustments were made to adhere to the eLife format.

