## Supplementary for "Longer mandible or nose? Co-evolution of feeding organs in early elephantiforms"

Supplementary Information

### Materials and methods

#### Materials

The materials examined in this work are from three longirostrine gomphothere families, i.e., Choerolophodontidae, Amebelodontidae, and “Gomphotheriidae” (*Gheerbrant and Tassy, 2009*). These materials include complete crania, mandibles, and teeth of different species, including *Choerolophodon chioticus*, *C. connexus*, and *C. guangheensis* (Choerolophodontidae); *Platybelodon dangheensis*, *Pl. grangeri*, *Pl. tetralophus*, *Pl. tongxinensis*, *Protanancus brevirostris*, *Protanancus* sp., *Pr. wimani* and *Aphanobelodon zhaoi* (Amebelodontidae); *Gomphotherium* cf. *angustidens*, *G. cooperi*, *G. inopinatum*, *G. steinheimense*, *G. tassyi* (“Gomphotheriidae”). All were housed in three museums: the Institute of Vertebrate Paleontology and Paleoanthropology (IVPP), American Museum of Natural History (AMNH), and Hezheng Paleozoological Museum (HPM; HMV is the specimen prefix). For the detailed specimen list, please see Data S2. These materials were discovered from four regions, the Linxia Basin, Tongxin region, Junggar Basin and Tunggur region (*Figures 2, S6*); these regions are fossil-rich, especially during the Shanwangian and Tunggurian stages (~20–11 Ma), with different fossil assemblages in different ages (*Deng et al., 2013*; *Wang et al., 2016*; *Wang et al., 2022*; *Qiu et al., 2013*). For the age information of each fossil assemblage (*Wang et al., 2022*; *Qiu et al., 2013*; *Sun, 2014*; *Wang, 2021*), please see Table S1.

#### Cladistic analysis

Cladistic analyses were performed to evaluate the phylogenetic hypothesis of trilophodont longirostrine proboscideans. The data matrix contained 37 taxa, including most of the known trilophodont longirostrine taxa at the species level, and *Phiomia serridens*, an Oligocene basal elephantiform, was selected as outgroup. Additionally, a basal elephantoid, *Tetralophodon longirostris* was also included to assess which clade the true elephantids originated from. The morphological characters included 5 characters from upper tusks, 9 from mandibular tusks, 37 from cheek teeth, 19 from the cranium, and 10 from the mandible, mainly following Tassy, 1996; Shoshani, 1996; and Wang et al., 2017. For a description of the characters and states, please see Appendix S1; for the data matrix, please see Data S1. Two methods, Bayesian tip-dating (BTD) and maximum parsimony (MP) analyses were performed.

In BTD analysis, the fossil ages were incorporated as tip calibrations (*Gavryushkina et al., 2014*; *Ronquist et al., 2012*; *Zhang et al., 2016*). The Lewis Mkv model (*Lewis, 2001*), with gamma rate variation across characters (Mkv + G) (*Yang, 1994*), was initially used; subsequently, the timetree was modelled by the fossilized birth death process (*Heath et al., 2014*; *Stadler, 2010*). The process was conducted using the time of the most recent common ancestor (root age) and included hyperparameters of speciation rate, extinction rate, fossil-sampling rate, and extant-sampling probability. The root age was first assigned an offset-exponential, with mean age of 37 Ma and minimum age of 34 Ma, referring to the oldest fossil. The fossil ages were fixed to their first occurrence. The extant-sampling probability was fixed to 1 because no living genera were specified. Apart from the timetree, the other key component was the relaxed clock model, which models the evolutionary rate variation along the branches in the tree. We used the independent gamma rate clock model (*Lepage et al., 2007*), in which the mean clock rate was initially assigned a lognormal prior (–6, 1) and the variance parameter of the clock rate was exponential (10).

We executed two independent runs and four chains per run (one cold chain and three hot chains) using Markov chain Monte Carlo. Each run was executed for 1 million generations and sampled every 5000 generations. The first 25% of the samples were discarded as burn-in and the rest of the two runs were combined. Good convergence and mixing were determined by effective sample sizes larger than 200 for all parameters and average standard deviations of split frequencies smaller than 0.01 (*Geyer, 1992*; *Ronquist et al., 2012a*). The BTD analysis was performed in MrBayes 3.2.7 (*Ronquist et al., 2012b*) (see Code Files S1).

MP reconstruction was performed by TNT1.1 (*Goloboff et al., 2008*). In MP analysis, all characters were equally weighted. Characters 20–25 pertained to the loph/lophid numbers of cheek teeth (character numbers begin from “1” in our numeration; however, in MP analysis, TNT1.1 automatically numbered characters from “0”; therefore, in the TNT1.1 program, these characters were numbered “19–24”). These were treated as ordered and irreversible, which was performed by setting “step-matrix → of costs” under the menu “Data → Character settings”, assigning the value “9” to the blanks where i > j (i to j in the matrix), and assigning the value “j–i” to the blanks where i < j. This setting was exported as the Code File S2, which can be re-loaded into the program. The traditional search strategy was performed, and the results were reported based on a 50% majority consensus tree of the most parsimonious trees. Node supports were calculated by symmetric resampling with 0.33 change probability (1000 replicates). The script files are provided in Code File S3. The major consensus tree was then calibrated using the timePaleoPhy function in R package paleotree 3.3.25 (*Bapst, 2012*), and is shown in Figure S1.

#### Stable carbon and oxygen isotope analysis

In this study, 83 tooth enamel samples from three gomphothere families were collected from gomphothere specimens (Data S3) for stable carbon and oxygen isotope analysis. We also compiled previously published isotope data of *Platybelodon* from the Tunggur region and Laogou, Linxia Basin (*Wang and Deng, 2005*; *Zhang et al., 2009*). The fossil teeth used in this study are well-preserved and showed no visible signs of alteration. Tooth enamel samples were obtained by cutting a small patch of enamel from a tooth or drilling along the entire length using a rotary drill; then, the samples were ground into fine powder. All samples (2–3 mg enamel powder) were then pre-treated with 5% sodium hypochlorite (NaOCl) overnight to remove any possible organic contaminants and then cleaned with distilled water. These samples were then treated with 1 M acetic acid overnight to remove non-structural carbonates and subsequently cleaned with distilled water. The treated samples were then freeze-dried.

The dried enamel samples were reacted with 100% phosphoric acid (H_3_PO_4_) at 25°C for approximately 72 hours. Carbon and oxygen isotope data were measured at Florida State University using a Finnigan MAT Scientific Delta Plus XP stable isotope ratio mass spectrometer coupled with a Thermo Scientific GasBench II. The lab standards that we used include MERK, MBCC, ROY-CC, and PDA. Results are reported in the standard delta (δ) notation as δ^13^C and δ^18^O values in reference to the international carbonate standard VPDB (Vienna Pee Dee Belemnite). We reconstructed the diet δ^13^C values of proboscideans from enamel δ^13^C values using an enrichment factor (ε*) of 13‰ for non-ruminants, like proboscideans (*Cerling and Harris, 1999*; *Passey et al., 2005*). Detailed results and specimen information are shown in Data S3.

#### FE analysis

We investigated the feeding behaviors of the three longirostrine gomphothere families, i.e., Choerolophodontidae, Amebelodontidae, and “Gomphotheriidae”, using finite element (FE) stimulation. Three species were selected to represent each family, *Choerolophodon chioticus*, IVPP V23457 (cranium and mandible); *Platybelodon grangeri*, HMV 0930 (cranium and mandible); and *Gomphotherium tassyi*, IVPP V22780 (cranium) and IVPP V22781 (mandible), respectively. Note that, in *Gomphotherium*, we were unable to get access to cranium and mandible that belonged to one individual. We used a handheld Artec Spider 3-dimensional scanner to obtain the surface topology of these specimens. The surface meshes were produced using Artec Studio 14 Professional. These meshes were first repaired with ZBrush 2021; for example, ZBrush 2021 was used to recover the broken edges of mandibular tusks, create the remaining mandibular tusks in the alveolus, create the keratinous cutting plate of the mandible in *Choerolophodon*, and for retro-deformation of the crushed cranium (i.e., the cranium of IVPP V23457).

The rough surface meshes were edited using Materialise 3-matic Research (V12.0). The surface meshes were smoothed by removing the small knobs and filling the holes for volume mesh generation. Before volume meshing, the models were cut into symmetric halves along the median sagittal plane. To reduce computation, only the right halves were preserved for further analyses. Note that the cranium was only used to define the attachments or insertions of jaw-closing muscles (modeled by many draglines, see below), and was treated as a rigid body in simulation (therefore, the cranial and nasal cavities are not relevant). The mandible contains two parts, the bony structure, including the cheek teeth, and the food acquisition organs (the mandibular tusk in *Platybelodon* and *Gomphotherium*, and the keratinous cutting plate in *Choerolophodon*). These two parts were integrated using the command “Non-manifold Assembly” in Materialise 3-matic Research. Then, the cranium and the mandible were aligned based on their natural position, i.e., the occlusal surface of the upper and lower tooth rows were matched, and the mandibular condyle and glenoid fossa were fitted. The sagittal surfaces of the cranium and mandible were made to coincide with the *x*-*z* plane, and the *x*-positive direction was set along the rostral direction. For comparison between different taxa, the three mandibular models were scaled to the same volume as *Choerolophodon* (6,563,708.0146 mm^3^) (*Figure S5*). Finally, volume meshes were generated in the cranium and mandible across the three models, and exported as .inp files that could be loaded into Abaqus CAE (V 6.14), the engineering software for FE analysis. The parameters for volume mesh generation are listed in Table S2.

The volume meshes, representing the geometric models, were imported into Abaqus CAE (V 6.14) (*Figures S6–S8*), and included two parts, cranium and mandible; additionally, a third part was created by Abaqus CAE, a long cylinder (300 mm in length, 50 mm in diameter), to model twigs that were cut by mandibular tusk or keratinous cutting plate. Note that the millimetre (mm)–ton (t)–second (s) unit system was adopted; other units included Newton (N) and million Pascal (MPa). Different materials, including bone, dentine, keratin, and wood were assigned to the corresponding parts. The materials of bone, dentine, and keratin were treated as isotropic linear elastic materials. For the detailed parameters, see Table S3 (*Drake et al., 2016; Huo et al., 2000*). However, twigs could not be treated as an isotropic linear elastic material, because the purpose of this simulation was to evaluate the food procuring efficiency of different taxa in different working conditions. Here the twigs were assigned as an orthotropic elastoplastic material, and the parameters (of wet red pine tree) were obtained from a wood handbook (*Risbrudt et al., 2010*) (Table S3).

The occlusal surfaces were coupled by two arbitrary points on the upper and lower teeth. These two points were connected by a “beam connector”, which constrained all the degrees of freedom (*df*) between the two points (*Figures S6A, S7A, S8A*). In this way, we simulated the occlusal surfaces of the upper and lower teeth. Jaw-closing muscles were simulated by several groups of “axis connectors” (*Fu et al., 2022*). This type of connector does not constrain any *df* of the two extreme points, and allows exerting force along the connector (*Figures* *S6B–E, S7B–E, S8B–E*). Four jaw-closing muscles were considered, including *temporalis* (*Figures* *S6B, S7B, S8B*), *superficial masseter* (*Figures* *S6C, S7C, S8C*), *zygomaticomandibularis* (*Figures* *S6D, S7D, S8D*), and *pterygoideus internus* (*Figures* *S6E, S7E, S8E*). Ten axis connectors were assigned to the *temporalis*, and four, three, and three were assigned to the latter three, respectively. These connectors were uniformly arranged along their insertion areas based on their natural anatomy. The areas of the temporal fossa (*At*) and ascending ramus (*AA*) were measured in 3-matic Research (V12.0) (*Figure S5*). We estimated the muscle force of *temporalis* as follows (*Tseng et al., 2017*):

*At* (mm^2^) × 0.3

this force was equally distributed to the 10 axis connectors for temporalis.

Alternatively,

*AA* (mm^2^) × 0.3

was considered the gross force for *superficial masseter*, *zygomaticomandibularis*, and *pterygoideus internus*. This force was also equally distributed to the other 10 axis connectors.

Note that the *At* and *AA* in the *Platybelodon* and *Gomphotherium* models were not true. These models were scaled to the same volume as that of *Choerolophodon*. In the simulation, we uniformly assigned muscle forces to make it easy to compare models (Table S4).

The cranium was treated as a rigid body and was fixed (*Figures* *S6A, S7A, S8A*). Another boundary condition was assigned to a node of the mandibular condyle, which only allowed the y-direction rotation and constrained any other *df*s (simulating the rotation of the mandibular articulation) (*Figures* *S6A, S7A, S8A*). The *df*s of the x- and z- rotations of the mid-symphysis were also constrained by considering the connection to the other half.

Two tests were carried out on the composite models: the distal forces test (*dft*) (Code S4) and twig-cutting test (*tct*) (Code S5). The *dft* includes two steps: 1, applying the muscle force; and 2, exerting a distal 5,000 N force that gradually changes from horizontally to vertically, by which we assessed the optimum direction of the external force for the mandible of each taxon. In this test, the “twig” was not included in the model.

In the *tct*, the middle point of the “twig” was set in close contact with the distal edge of the mandibular tusk and keratinous cutting plate (*Figures* *S6A, S7A, S8A*), and was placed horizontally (Figures S6), 45° obliquely (*Figures S7*), and vertically (*Figures S8*). One extremity of the “twig” was fixed, and contact properties were assigned (hard contact normally and 0.3 frictional coefficient tangentially). The *tct* also includes two steps: 1, applying the muscle force as in *dft*; and 2, displacing the cranium and mandible 10 mm towards the “twig” to stimulate cutting action of proboscideans, by which we determined the cutting efficiency of directions for each taxon. In the results, the sum of the equivalent plastic strain (EPS) from total twig elements was calculated and reported for each model. The plastic strain represents the irreversible deformation of an element, and the sum of EPS from all twig elements can reflect the cutting effects in each model. The movies for von Mises stress contour colour maps were also generated in the *dft* (Movies S1–S3) and *tct* (Movies S4, S6, S8, S10, S12, S14, S16, S18) modelling, and for the EPS contour colour maps of twigs in *tct* (Movies S5, S7, S11, S13, S15, S17, S19) modelling.

#### Principal components analysis (PCA) and PC scores mapping on the tree

In PCA, only taxa of the three gomphothere families—members of Choerolophodontidae, Amebelodontidae, and “Gomphotheriidae”, in addition to *Phiomia serridens*—were retained; the mammutid taxa (*Losodokodon*, *Eozygodon*, and *Zygolophodon*) and the stem elephantimorphs (*Eritreum* and *Gomphotherium annectens*) were excluded from the analyses.

A character combine, including characters 1–14 of upper and lower tusks, as well as 71–73, 76, 77, and 80 of the mandibles (Supplementary Appendix S1 and Data S1), was generated using PCA, which represents the synthetic character states of food acquisition organs. Besides, another two character-combines, four characters concerning the narial region (i.e., characters 54–57; see Supplementary Appendix S1 and Data S1) and five characters (characters 5, 9, 11, 72, and 77) in relation to the horizontal cutting behavior, were also generated using PCA. PCA was performed in Past 4.04, in which the missing values were predicted using mean value imputation. The data on PC1 vs. PC3 or PC2 vs PC3 plans were plotted (*Figure S3A, C, D*).

The PC1 of the food acquisition organ combination (*Figure S3A*) and that of the narial region (*Figure S3C*), as well as the PC2 of the character-combine in relation to the horizontal cutting behavior (*Figure S3D*) were selected to represent the synthetic evolutionary state of each. We use PC2 rather than PC1 in the last test because the characters in relation to the horizontal cutting behavior were not evolved in one-way. For example, the character 9 indicates the width of the mandibular tusks, which includes three states: 0, wide; 1, very wide; 2, narrow. In this case, the larger value (2 = narrow mandibular tusker) does not mean the stronger horizontal cutting effect. Finally, these PCs were respectively mapped on the BTD tree (mammutids and stem elephantimorphs removed), using the contMap function in R package phytools 0.7-90 (*Bapst, 2012*).

### Supplementary results and discussion

#### Phylogenetic reconstruction

To comprehensively explore the phylogenetic relationships among trilophodont longirostrine proboscideans, especially for the longirostrine bunodont elephantiforms, BTD and MP analysis were performed.

**BTD** (*Figure 1A*): The resulting tree had two major branches at the base, i.e., mammutids (100% node support) and bunodont elephantiforms (91% node support), which are represented by zygodont and bunodont cheek tooth morphology. In the bunodont elephantiforms, *Eritreum* and *Gomphotherium annectens* were stem taxa. *Gomphotherium annectens* did not cluster with the other members of “Gomphotheriidae”; this supports the assertion of Tassy that the “*G. annectens*” group is not closely related to other bunodont elephantiforms (*Tassy, 1994*). The other in-group taxa constitute three monophyletic groups: Choerolophodontidae (100% node support), Amebelodontidae (99% node support), and “Gomphotheriidae” + *Tetralophodon longirostris* (59% node support), with the last taxon representing the root of Elephantoidea. Therefore, “Gomphotheriidae” (except for the “*G. annectens*” group) might be paraphyletic.

In Choerolophodontidae, because *Gnathabelodon throperi* was inserted among *Choerolophodon* species, *Choerolophodon* will be further studied by us (*Li et al., 2019*). In Amebelodontidae, *Platybelodon* and *Aphanobelodon* species constituted a monophyletic group characterized by a very wide mandibular symphysis and tusks (*Wang et al., 2017*); members of this group represent typical “trunk grasping” and mandibular tusk cutting taxa within amebelodontids. Other groups represent generalists in Amebelodontidae. It is notable that *Megabelodon* and *Eubelodon* (*Barbour, 1932; Barbour, 1934*), although lacking mandibular tusks, are deeply embedded within Amebelodontidae because of the high similarity of check teeth with some amebelodontid taxa, especially for *Protanancus* *wimani* (*Wang et al., 2015*). For “Gomphotheriidae”, two groups were also derived (*Wang et al., 2017*) One is the sub-bunodont group, which have a slight zygodont character in their cheek teeth. *Rhynchotherium* was also included in this group, which might have further evolved to the brevirostrine *Cuvieronius* (*Mothé et al., 2016*). The other group is characterized by highly bunodont cheek tooth morphology, and *Tetralophodon longirostris*, perhaps with other true elephantoids, was derived from this clade (*Wu et al., 2018*).

**MP analysis** (*Figure S1*): The 36 most parsimonious trees of 253 steps resulted in a CI of 0.470 and RI of 0.779. A 50% majority strict consensus tree is shown here, which was similar to the BTD tree. The two major groups were still present, with 100% node support for Mammutidae, and 45% node support for bunodont elephantiforms. Similar to BTD, *Eritreum* and *Gomphotherium annectens* are positioned as the stem taxa of bunodont elephantiforms, but *G. annectens* did not cluster with the other *Gomphotherium*. Choerolophodontidae (97% node support) were monophyletic and differentiated after *Eritreum* and *G. annectens*. Amebelodontidae (46% node support) was another monophyletic group that derived from the paraphyletic “Gomphotheriidae” (59% node support). Within Amebelodontidae, *Serbelodon*, *Archaeobelodon*, and two species of Chinese “*Protanancus*” (*Pr. brevirostris* and *Pr. wimani*) were stem groups. *Platybelodon* + *Aphanobelodon*, the “trunk grasping” and mandibular tusk cutting taxa constituted one monophyletic group, and the other taxa constituted another monophyletic group, including the tuskless taxa *Megabelodon* and *Eubelodon*. The clade of sub-bunodont *Gomphotherium* (and *Rhyncotherium*) was positioned as the sister group of Amebelodontidae, and they together further constituted a sister group of “highly bunodont *Gomphotherium*” + *Tetralophodon longirostris*, which probably led to the evolution of elephantoids.

In both BTD and MP analyses, Choerolophodontidae and Amebelodontidae were monophyletic groups with distinct mandible and mandibular tusk morphology. Choerolophodontidae lost their tusks but became equipped with a keratinous cutting plate, which was highly efficient for vertically cutting vegetation. Amebelodontidae developed wide mandibular tusks (except for *Megabelodon* and *Eubelodon*), and, particularly in the sub-group “*Platybelodon* clade”, the mandibular tusks became extremely wide with a distal cutting edge, which was specialized for horizontally cutting vegetation. In both analyses, “Gomphotheriidae” were not monophyletic. However, all members of “Gomphotheriidae” possessed clubbed mandibular tusks and narrow symphysis. This morphology is rather unspecialized, which indicates rather general feeding habitats. It should be noted that many later taxa, such as true elephantoids and possibly some “brevirostrine bunodont elephantiforms”, were derived from “Gomphotheriidae”.

#### Comparative morphology of narial region and mandibular symphysis among *Platybelodon*, *Gomphotherium*, and *Choerolophodon*.

In longirostrine bunodont elephantiforms, the mandibular symphysis with mandibular tusks (or keratinous cutting plate) was the primary feeding organ, and the developing trunk was the auxiliary feeding organ. The development of a trunk can be deduced by the morphology of the narial region.

**Mandibular symphysis**: In all longirostrine bunodont elephantiforms, the mandibular symphysis was obviously elongated, however, to a different extent (*Gheerbrant, and Tassy, 2009*). They also differed in thickness, distal expansion, and concaveness, depending on the morphology of the mandibular tusks which it holds.

*Gomphotherium* possesses the least specialized mandibular symphysis and mandibular tusks (*Figures 1K, S2B*). The symphysis is the shortest and narrowest among the three groups. The distal part is only slightly transversely expanded. A narrow and moderately deep groove is developed on the dorsal surface of the symphysis, between the two alveolar crests. In lateral view, the symphysis is the thickest among the three groups. There are three or four mental foramina on each side of the symphysis and the following mandibular corpus. The rostral-most one is the largest, but is only slightly larger than the others. Each mandibular tusk is long-rod like. The tusk is slightly twisted. A conspicuous dorsal groove is along the dorsal surface, and another weak ventral groove is also present. The distal end tapers to the middle axis due to wear. An elongated, rounded triangular wear facet is very clear. The ventral surface is also polished. The cross-section at the alveolus is obliquely pyriform, with a typical concentric lamellar structure.

*Platybelodon* possesses the largest mandibular symphysis among the three groups (*Figures 1I, J, S2A*). The symphysis is the longest and the distal part is most expanded. The symphysis is deeply dorsally concave, bordered by two conspicuous alveolar crests. Because of the concaveness, the symphysis is strongly pocket-like in lateral view. The proximal end has a strong transverse ledge, and the distal end (openings of the alveoli) is nearly straight. Each mandibular tusk is extremely flattened and plate-like. The tusk is also slightly twisted; the tusk cross-section is nearly vertical at the proximal end, becomes horizontal distally, and there are dentinal rods throughout the section. In dorsal and ventral views, the distal end is straight, regardless of the small breaks; and in distal view, it is sharpened ventrally with a cutting edge, like a wedge. Additionally, the strongly cutting wedge morphology of mandibular tusks is only present in the “*Platybelodon* lineage”. In the other members of Amebelodontidae, although the mandibular tusks (if present) are more or less flattened, and the distal end of the symphysis is expanded, and the cutting-edge morphology is not pronounced (*Lambert, 1992*).

The mandibular symphysis of *Choerolophodon* *chioticus*, possibly in all choerolophodontids, is the most specialized among the three groups, in terms of lacking mandibular tusks (*Konidaris et al., 2016*; *Li et al., 2019*) and development of the keratinous cutting plate (*Figures 1L, M, S2C–F*). The symphysis is moderately long, with a slightly distal expansion. It is wider than that of *Gomphotherium*, but narrower than that of *Platybelodon*. The symphysis is strongly concave and trough-like. The bony wall is very thin because of the complete absorption of tusk alveoli. Two or three mental foramina are present on each side, and the rostral one is extremely large. It is tube-like, which indicates strong development of mental nerves and the eponymous artery for nourishing the keratinous cutting plate that is of dermal origin (*Sisson, 1953*) (*Figures* *S2D*). There is a very deep and narrow slit present on each side of the distal alveolar crest (or distal mandibular trough edge), which is presumably for holding the keratinous cutting plate (*Figures 1L, S2C–F*). Because the holding slit is bent along the bending bony sheath, the keratinous cutting plate may have a vertical dorsal cutting edge, to balance growth and wear (see the reconstruction, *Figures* *1M*). The normal position of the tusk alveolus was vestige, showing a rough and slightly depressed area. This morphology is very similar to that of *Megabelodon lulli*, which also lacks mandibular tusks (*Barbour, 1934*). The dorsal and ventral surfaces of the distal symphysis is rough with numerous vascular pores; this further indicates the presence of keratinous integument, perhaps to protect the distal symphysis.

**Narial region**: The evolutionary level of the trunk can be completely inferred from the morphology of the narial region (*Tassy, 1994*). Here we first showed the narial region of a living elephant (*Elephas maximus*, IVPP OV733) (*Figure 1E*), focusing on the following four morphological factors. 1, the dorsal border of the narial aperture is slightly caudal to the postorbital process. This part is for the attachment of *maxilla-labialis*, which is the key muscle for manipulating the entire trunk (*Boas and Paulli, 1908*; *Eales, 1926*). 2, the narial aperture is wide, showing a pair of deep and sub-circular perinasal fossae. This part related to the insertion of *lateralis nasi*, and functions in enlarging the nostril cavity to suck up water in the trunk (*Boas and Paulli, 1908*; *Eales, 1926*). 3, the nasal process of the nasal bone is moderately developed. 4, the insertion slit for the mesethmoid cartilage is narrow and small, and deeply concealed in the narial aperture. The latter two points might be related to trunk flexibility. The smaller the nasal bone process and mesethmoid cartilage, the more flexible the trunk. These points were carefully discussed by Tassy (*Shoshani et al., 2006*).

*Gomphotherium tassyi*, which is phylogenetically closer to living elephants, also shows similar narial aperture morphology (*Figures 1G, S9D, E*). The dorsal border of the narial aperture is more caudally positioned than that of *Elephas*, as it is somewhat distantly caudal to the postorbital process. The perinasal fossa is dorso-ventrally narrower and shallower than that of *Elephas*. It is also latero-ventrally oblique compared with that of *Elephas*. The nasal process of the nasal bone is comparable to that of *Elephas*. The slit for mesethmoid cartilage insertion is somewhat wider than that of *Elephas*, and is still deeply within the narial aperture. In general, the evolutionary level of the narial region of *G. tassyi* is comparable to that of *Elephas*, but maybe slightly more primitive. Furthermore, the morphology of the narial region of *G. tassyi* is also similar to some other *Gomphotherium* species, such as *G. angustidens* and *G. productum* (*Tassy, 2013*; *Osborn, 1936*), however, it is more derived than that of *G. annectens*, which might not be phylogenetically close to other *Gomphotherium* species, see Tassy, 1994 and the phylogeny in Figure 1A.

*Choerolophodon* shows a relatively more primitive evolutionary stage of narial aperture morphology (*Figures 1H, S9A–C*), as Tassy, 1994 stated. In our choerolophodontid sample, the dorsal border of the narial aperture is also slightly caudal to the postorbital process, like in *Elaphas*. In *C. chioticus*, although the narial aperture has been enlarged, the perinasal fossa is only weakly developed, presenting like a dorso-ventrally narrow groove (red area in *Figures* *1H*); however, in the more primitive *C. guangheensis*, the perinasal fossa is totally absent (*Figures* *S9A–C*). The nasal process of the nasal bone is very large. It is nearly rectangular rather than triangular. The slit for mesethmoid cartilage insertion is wide and large, which can be clearly seen from outside. The morphology of the *Choerolophodon* narial region was clearly primitive and indicated a weakly developed trunk compared with living elephants and *Gomphotherium*.

Tassy, 1994, also showed a comparable evolutionary stage of narial region in an amebelodontid *Archaeobelodon filholi*, but did not mention *Platybelodon*. Indeed, the narial region of *Platybelodon* was not carefully studied until Wang and Li, 2022. In *Platybelodon*, the narial region is very specialized (*Figures 1F, S9G–L*). The dorsal border of the narial aperture is distantly caudal to the postorbital process, like in *Gomphotherium*, but more importantly, the narial aperture is greatly enlarged; this provides a very large area for attaching *maxilla-labialis*, the most important part of trunk muscle group. Apart from the latero-ventrally oblique perinasal fossa that it similar to that in *Gomphotherium*, on each premaxilla, a vast inclined region is rostral to the perinasal fossa, called the prenasal slope, which possibly provides additional attachment for *nasialis* (*Boas* *and Paulli, 1908*). Furthermore, the nasal process of the nasal bone is very small, only slightly rostrally and dorsally bulged from the apertural dorsal border, and the insertion for mesethmoid cartilage is very small, which indicates the very weakly developed mesethmoid cartilage.

Additionally, we analysed a large sample of *Platybelodon* *grangeri* crania from the Linxia Basin housed in HPM, and the observed features were all clear in adult individuals (*Wang et al., 2013*) (*Figures 1F, S9G–L*). In the other *Platybelodon* species, e.g., in *Pl. tongxinensis* and *Pl. tetralophus*, these features were also generally developed (*Wang and Li, 2022*) (*Figures S9K, L*), and were even found in *Aphanobelodon zhaoi* (*Wang et al., 2017*) (*Figures S9F*). These taxa constitute a monophyletic lineage within Amebelodontidae (*Figures 1, S1*). However, in the other taxa, such as *Protanancus* spp., which was replaced with *Platybelodon* in northern China, the specialized morphology of the narial region is absent (*Wang et al., 2015*). In four critical morphological factors for trunk development, *Platybelodon* is even more derived than living *Elephas*, which indicates a very developed trunk in *Platybelodon* that could perform some actions like living elephants (*Purkart et al., 2022*). However, *Platybelodon*, along with the other longirostrine bunodont elephantiforms, have two infraorbital foramina, whereas elephantoids have a very large infraorbital foramen (*Tassy, 2013*); this possibly indicates a less tactile trunk relative to the enlargement of the maxillary branch of the trigeminal nerve (Westerhold et al., 2020).

#### PCA

PCA was respectively performed for the characters of the narial region and the food acquisition organs (mandible and tusks) to extract their synthetic characters.

For the food acquisition organs (*Figures* *S3A*), the PC1–PC3 plan also shows a good distribution of all taxa. *Phiomia* positions itself at the lower-left corner, along with several other taxa, including some sub-bunodont *Gomphotherium* species and some stem amebelodontids (*Protanancus brevirostris*, *Archaeobelodon filholi*, et al.). Choerolophodontidae are located on the right side of the plan, away from the other taxa, and have large sores of PC1 (mostly loaded by upper tusk characters). *Platybelodon*, being at the top of the plan, has large scores of PC3, which are heavily loaded by longer and wider mandibular symphyses (C71 and C72 in the plot).

For the narial region (*Figures* *S3C*), the data points were largely overlapped, because of the relatively few character states and, partially, because of the missing data. In the PC1–PC3 plan, *Phiomia* is located at the lower-left corner, far from the other data points. The character 54 (position of the nasal aperture, C54 in the plot), contributes the most to PC3 and greatly separates *Phiomia* from the other taxa. PC1 appears to be a good representation of the evolutionary level of the narial region, with *Platybelodon* and *Aphanobelodon* having the largest sores and *Phiomia* having the smallest. Choerolophodontidae usually have smaller PC1 scores than other taxa, indicating the relatively primitive evolutionary level of the narial region.

For the characters in relation to the horizontal cutting behavior (*Figures S3D*), we found that PC2–PC3 plan also shows a well separation of different taxa, except taxa of Choerolophodontidae (all clustered at the upper-left corner), in which the mandibular tusks were lost, and therefore shows little difference in horizontal cutting behavior. Taxa of *Platybelodon* clade were all at the right part of the plan, indicating the stronger behavior of horizontal cutting. Taxa of *Gomphotherium* all situate at the lower-left part with some amebelodontid taxa, and other amebelodontid taxa, like *Protanancus macinessi*, and *Pr. chinjiensis*, were in the middle of the plan.

For presenting the evolutionary level of the above character-combines in different lineages, the above PC1s or PC2 were mapped on the BTD phylogenetic tree, in which mammutids and stem elephantimorphs have been removed. For PC1 of tusks and mandible combine (*Figures S3B*), the blue colors of Choerolophodontidae suggest the very aberrant and specialized mandibular and tusk morphology, as a compensation of the less developed trunk (see below). This PC1 values of Choerolophodontidae are followed by the *Platybelodon* lineage, marked by the green colors, also revealing their speciation of mandible and tusks. This character-combine are relatively unspecialized in other taxa.

For PC1 of the narial region (*Figure 4A*), the pure blue color of *Platybelodon* (except *Pl. barnumbrowni* from North America, of which the color is closer to green rather than blue) and *Aphanobelodon* reveals the highly developed narial region of this lineage, while Choerolophodontidae show red colors, indicating their relatively low evolutionary level of narial region. The other taxa show moderately evolutionary level narial region. Interestedly, the evolutionary level of the character-combine in relation to horizontal cutting is highly correlated with that of narial region (*Figure 4B*). In the *Platybelodon* clade, this character-combine is also in pure blue color (except *Pl. barnumbrowni*); while in the Choerolophodontidae clade, the red or yellow colors indicate relatively low evolutionary levels, which is also comparable with that of narial region. Colors in the other taxa also show from slow to moderately evolutionary level. The result strongly indicates a highly co-evolution between narial region and horizontal cutting behavior in trilophodont bunodont elephantiforms.

#### Geographic distribution and relative abundance of various bunodont elephantiforms

Fossil bunodont elephantiforms were very abundant from the Early to Middle Miocene strata of northern China, ranging from ~19–11.5 Ma, and mainly in the following regions: the Linxia Basin, Tongxin region, Junggar Basin, and Tunggur region (*Deng et al., 2013*; *Wang et al., 2016*; *Wang et al., 2022*; *Qiu et al., 2013*) (*Figure* *S10*). In each region, bunodont elephantiforms were distributed into different fossil assemblages based on age (*Wang et al., 2022*; *Qiu et al., 2013*; *Sun, 2014*; *Wang, 2021*) (*Figure 2, Table S1*) and components. See details below:

The Linxia Basin was subdivided into four assemblages: Dalanggou Fauna (LX1, ~19–18 Ma), including *C. guangheensis* (Choerolophodontidae), *Pr. brevirostris* (Amebelodontidae), *G. inopinatum*, and *G.* cf. *angustidens* (“Gomphotheriidae”); Sinanu Fauna (LX2, ~17–15.2 Ma), including *Pr. wimani*, *Pl. tongxinensis* (Amebelodontidae), *G. inopinatum*, and *G. tassyi* (“Gomphotheriidae”); Zengjia Fauna (LX3, ~14.5–14 Ma), including *Pl. grangeri* (Amebelodontidae) and *G. tassyi* (“Gomphotheriidae”); and the Laogou Fauna (LX4, ~14–12.5 Ma), which had the same gomphothere components as the Zengjia Fauna.

The Tongxin (and Zhongning) region also had four assemblages: Lower Miaoerling (A/B zones) Fauna (TX1, ~17.8–17 Ma), including *C. chioticus* (Choerolophodontidae), *Pr. wimani* (Amebelodontidae), *G. inopinatum*, and *G.* *cooperi* (“Gomphotheriidae”); Middle Miaoerling (C zone) Fauna (TX2, ~16.6–16 Ma), including *C. chioticus* (Choerolophodontidae), *Pr. wimani*, and *Pl. tongxinensis* (Amebelodontidae); Upper Miaoerling (D/E) Fauna (TX3, ~14.5–13.5 Ma, = Maerzuizigou Fauna in Dingjiaergou), including *Aphanobelodon zhaoi*, *Pl. grangeri* (Amebelodontidae), and *G. tassyi* (“Gomphotheriidae”); and Heijiagou Fauna (TX4, ~13.4–12.7 Ma), including *Pl. grangeri* (Amebelodontidae) and *G. tassyi* (“Gomphotheriidae”).

The northern Junggar Basin also had four assemblages: Top Suosuo Fauna (JG1, ~18.8–18.4 Ma), only including *Protanancus* sp*.*; Halamagai Fauna (JG2, ~16.9–15 Ma), including *C. connexus* (Choerolophodontidae), *Protanancus* sp*.*, *Pl. dangheensis* (Amebelodontidae), *G. tassyi*, and *G. steinheimense* (“Gomphotheriidae”); Kekemaideng Fauna (JG3, ~14.5–14 Ma), including *C. connexus* (Choerolophodontidae), *Pl. grangeri* (Amebelodontidae), and *G. tassyi* (“Gomphotheriidae”); and Dingshanyanchi I Fauna (JG4, ~13–12.6 Ma), which was primarily characterized by small mammals and only included *Pl. grangeri* (Amebelodontidae).

The Tunggur region included three successive assemblages: Tairum (TG1, ~13.6–13.2 Ma), Moergen (TG2, ~13.2–12 Ma), and Tamuqin (TG3, ~12–11.6 Ma). Among the faunas, only one genus, *Platybelodon*, was found, with *Pl. grangeri* from the Tairum and Moergen Faunas, and *Pl. tetralophus* from the Tamuqin Fauna.

Based on the gomphothere taxa secession of the major regions of northern China, we found that bunodont elephantiforms primarily occurred after ~19 Ma and ended ~11.5 Ma. The three families, Choerolophodontidae, “Gomphotheriidae”, and Amebelodontidae, were all present, at the beginning represented by *Choerolophodon*, *Protanancus*, and *Gomphotherium*, respectively. During ~17–15 Ma, i.e., the Mid-Miocene Climate Optimum (MMCO), *Platybelodon* appeared first, and was gradually replaced by *Protanancus*. After ~14.5 Ma, i.e., the beginning of the Mid-Miocene Climate Transition (MMCT), *Choerolophodon* last appeared in northern China. After ~12.5 Ma, *Gomphotherium* were also extinct and only *Platybelodon* existed. However, mammutids continuously persisted until ~11.5 Ma.

To approximately quantify the relative abundance of each gomphothere taxon in different ages, we summarized the specimens housed in IVPP, HPM, and AMNH, and calculated the percentage of each taxon in different faunas (*Figures 2, S4*). Of the Dalanggou Fauna (LX1), *C. guangheensis* represented 35.5%; *Pr. brevirostris*, 60.5%; and *Gomphotherium*, 3.9%. Of the Lower Miaoerling Fauna (TX1), *Choerolophodon chioticus* represented 27.3%; *Pr. wimani*, 54.5%; *Gomphotherium*, 13.6%; and Mammutidae, 4.6%. Of the Halamagai Fauna (JG2), *C. connexus* represented 35.5%; *Protanancus* sp., 1.6%; *Pl. dangheensis*, 25.8%; *Gomphotherium*, 19.4%; and Mammutidae, 17.7%. Of the Kekemaideng Fauna (JG3), *C. connexus* represented 7.1%; *Pl. grangeri*, 64.3%; and *G. tassyi*, 28.6%. Of the Upper Miaoerling Fauna (TX3), *A. zhaoi* represented 40%; *Pl. grangeri*, 44%; *G. tassyi*, 12%; and Mammutidae, 4%. Of the Zengjia + Laogou Fauna (LX3,4), *Pl. grangeri* represented 79%; *G. tassyi*, 20%; and Mammutidae, 1%. Of the Hejiagou Fauna (TX4), *Pl. grangeri* represented 80% and *G. tassyi* represented 20%. Finally, of the Tunggur region (all three faunas, TG1–3), *Platybelodon* represented 97.2%, and Mammutidae represented 2.8%.

Based on the above data, before ~17 Ma, *Choerolophodon* and *Protanancus* were the predominant proboscideans; *Gomphotherium* and Mammutids were rare. During ~17–15 Ma, i.e., the MMCO, *Choerolophodon* and Mammutids reached their highest relative abundance, with *Protanancus* being replaced with *Platybelodon*. After ~14.5 Ma, the beginning of the MMCT, *Platybelodon* significantly increased in the faunas. *Choerolophodon* tended to be extinct and the relative abundance of *Gomphotherium* also increased. After ~12.5 Ma, *Gomphotherium* was also extinct, and *Platybelodon* became the predominant proboscidean and the mammutids survived until the end of the Middle Miocene. Therefore, the beginning of the MMCT was accompanied by the rise and thriving of *Platybelodon*, when the open environment was dominated by grasslands due to the prevalent relative aridity found in northern China.

**Supplementary Appendix**

Appendix S1. Morphological characters in the phylogenetic analyses

Upper tusk

1. Thickness, in adult male: thick (0); thin (1); absent (2); very thick (3).
2. Bend: ventrally bent (0); straight (1); dorsally bent (2).
3. Spirality: not spiral (0); slightly spiral (1); strongly spiral (2).
4. Diverge of the two tusks: moderately diverge (0); parallel (1); strongly diverge (2).
5. Enamel: with an enamel band (0); without enamel (1); enamel enclosing the entirely tusk (2).

Lower tusk

6. Presence: present (0); absent (1).
7. Bend: dorsally bent (0); straight (1).
8. Spirality: slightly spiral (0); straight (1).
9. Width: wide (0); very wide (1); narrow (2).
10. Enamel: without enamel (0); with an enamel band (1).
11. Applanation of cross-section: flattened (0); extremely flattened (1); pyriform (2); round (3).
12. Median edge: straight (0); round (1).
13. Structure of cross-section: concentric laminae (0); dentine rods (1).
14. Length beyond alveolus: long (0); short (1).

Cheek teeth

15. DP2: possessing a large anterior cone (0); possessing a weak anterior cone (1).
16. DP3 and dp3: possessing only two lophs/lophids (0); possessing two lophs/lophids plus a strong posterior cingulum/cingulid (1); possessing three lophs/lophids (2).
17. DP3 and dp3, second loph/lophid: straight (0); oblique (1).
18. DP3: cones aligned (0); cones alternatively arranged (1).
19. dp3: conids aligned (0); conids alternatively arranged (1).
20. Dp4/dp4: possessing two lophs/lophids plus a strong posterior cingulum/cingulid (0); possessing three lophs/lophids (1); possessing four lophs/lophids (2).
21. Premolars: p2 present (0); p3 present (1); p4 present (2); p4 absent (3).
22. M1/m1: trilophodont (0), tetralophodont (1).
23. M2/m2: trilophodont (0), tetralophodont (1).
24. M3: 3^rd^ loph incomplete (0); 3^rd^ loph complete (1); 4^th^ loph incomplete (2); 4^th^ loph complete (3); pantalophodont (4).
25. m3: trilophodont (0); 4^th^ lophid incomplete (1); 4^th^ lophid complete (2); pantalophodont (3); hexalophodont (4).
26. upper molars, the 2^nd^ loph: posterior pretrite central conule weak or absent (0); posterior pretrite central conule well-developed (1).
27. upper molars, the 2^nd^ loph: posterior pretrite central conule smaller than, at most equivalent to the anterior pretrite central conule (0); posterior pretrite central conule larger than the anterior pretrite central conule.
28. lower molars, the 2^nd^ lophid: anterior pretrite central conule weak or absent (0); anterior pretrite central conule well-developed (1).
29. lower molars, the 2^nd^ lophid: posterior pretrite central conule not enlarged (0); posterior pretrite central conule enlarged (1).
30. molars, typically in the 2^nd^ loph/lophid: crescentoids absent (0); crescentoids weak (1); crescentoids complete (2).
31. molars, typically in the 2^nd^ loph/lophid: posttrite mesoconelet(s) well-developed (0); posttrite mesoconelet(s) weak or absent (1).
32. molars, typically in the 2^nd^ loph/lophid: pretrite mesoconelet individualized (0); pretrite mesoconelet fused with the anterior pretrite central conule (1).
33. molars, typically in the 2^nd^ loph/lophid: posttrite central conule(s) absent (0); posttrite central conule(s) present (1).
34. upper molars, typically in the 2^nd^ loph: posttrite half loph divided into two to three elements (main conelet and mesoconelet(s)) (0); posttrite half loph subdivided more than three elements (2); posttrite half loph subdivided as a thin crest (3).
35. molars: elements of posttrite half lophs/lophids inflated (0); posttrite half lophs/lophids compressed (1); posttrite half lophs/lophids highly compressed as a thin crest (2).
36. molars, typically in the 2^nd^ loph/lophid: each pretrite central conule composed of 1~2 conules (0); each pretrite central conule composed of more than 3 conules, showing a thick crest (1); pretrite central conule absent (2).
37. upper molars, zygodont crest: absent (0); thick (1); thin (2).
38. lower molars, zygodont crest: absent (0); present (1).
39. molars, typically in the 2^nd^ loph/lophid: pretrite trefoil incomplete (0); pretrite trefoil well-developed (1); pretrite trefoil with thin, crest-like lobes (2); pretrite trefoil secondary weakened or at least showing the tendency (3).
40. molars, typically in the 2^nd^ loph/lophid: posttrite trefoil absent (0); posttrite trefoil incomplete (1); posttrite trefoil complete (2).
41. upper molars, typically in the 2^nd^ loph: pretrite and posttrite half lophs aligned (0); pretrite and posttrite half lophs more or less chevroned (1).
42. lower molars, typically in the 2^nd^ lophid: pretrite and posttrite half lophids aligned (0); pretrite and posttrite half lophids more or less chevroned (1).
43. lower molars, typically in the 2^nd^ lophid: posttrite half lophid normal to mid-axis (0); posttrite half lophid distadaxially oblique (1).
44. molars: without anancoid contact (0); with pseudanancoid contact (1).
45. upper molars, typically in the 1^st^ interloph: entoflexus “I-shaped” (compressed) (0); entoflexus “V-shaped” (1); entoflexus open (2).
46. lower molars, typically in the 1^st^ interlophid: ectoflexid “V-shaped” (compressed) (0); ectoflexid “U-shaped” (1); ectoflexid open (2); ectoflexid “I-shaped” (highly compressed) (3).
47. molars, width: in normal width (0); narrow (1); broad (2).
48. molars, ptychodonty: absent (0); present (1).
49. molars, choerodonty: absent (0); present (1).
50. molars, cementodonty: absent (0); present (1).
51. molars, crown height: low crowned (0); slightly high crowned (1).

Cranium

52. braincase: narrow (0); wide (1).
53. braincase: low (0); relatively high (1).
54. narial, nasal aperture: rostral to the orbit (0); upon the orbit (1).
55. narial, perinasal fossa: absent or weak (0); with a complete perinasal fossa (1); extremely large, showing a prenasal slope (2).
56. nasal bone, nasal process: large (0); small (1).
57. mesethmoid cartilage insertion: large (0); small (1).
58. subnasal fossa: absent (0); present (having an incisive constriction) (1).
59. rostrum: narrow (0); wide (1).
60. rostrum: short (0); long (1).
61. lachrymal opening: present (0); absent (1).
62. orbital: in normal position (0); caudally positioned (1); dorsally positioned (2); rostrally positioned (1).
63. orbitotemporal crest: oblique (0); vertical (1).
64. facial region: rostrally positioned (0) caudally positioned (1); ventrally stretched (2).
65. post palatine: with a spine (0); with a tuberosity (1); without ornamentation (2).
66. basicranium: not erected (0); erected (1).
67. zygomatic processes: wide in distance (0); narrow in distance (1).
68. tympanic: narrow (0); inflated (1).
69. tympanic: separated foramen lacerum medium and foramen ovale (1); merged foramen of the above.
70. tympanic: internal carotid artery foramen large (0); internal carotid artery foramen small or absent (1).

Mandible

71. symphysis: long (0); short (1); extremely elongated (2).
72. symphysis: wide (0); narrow (1); extremely widened (2).
73. symphysis: shallow (0); deep (1).
74. symphysis: with keratinous structure (0); without keratinous structure (1).
75. symphysis: with a transverse ledge at the proximal end (0); without a transverse ledge at the proximal end (1).
76. symphysis: not ventrally bent (0); ventrally bent (1).
77. symphysis: proximal end close to the tooth row (0); that distant from the tooth row (1).
78. angular process: not elevated (0); elevated (1).
79. angular process: posteriorly protruded (0); not posteriorly protruded (1).
80. ramus: vertical (0); caudally inclined (1).

#### Supplementary Figures

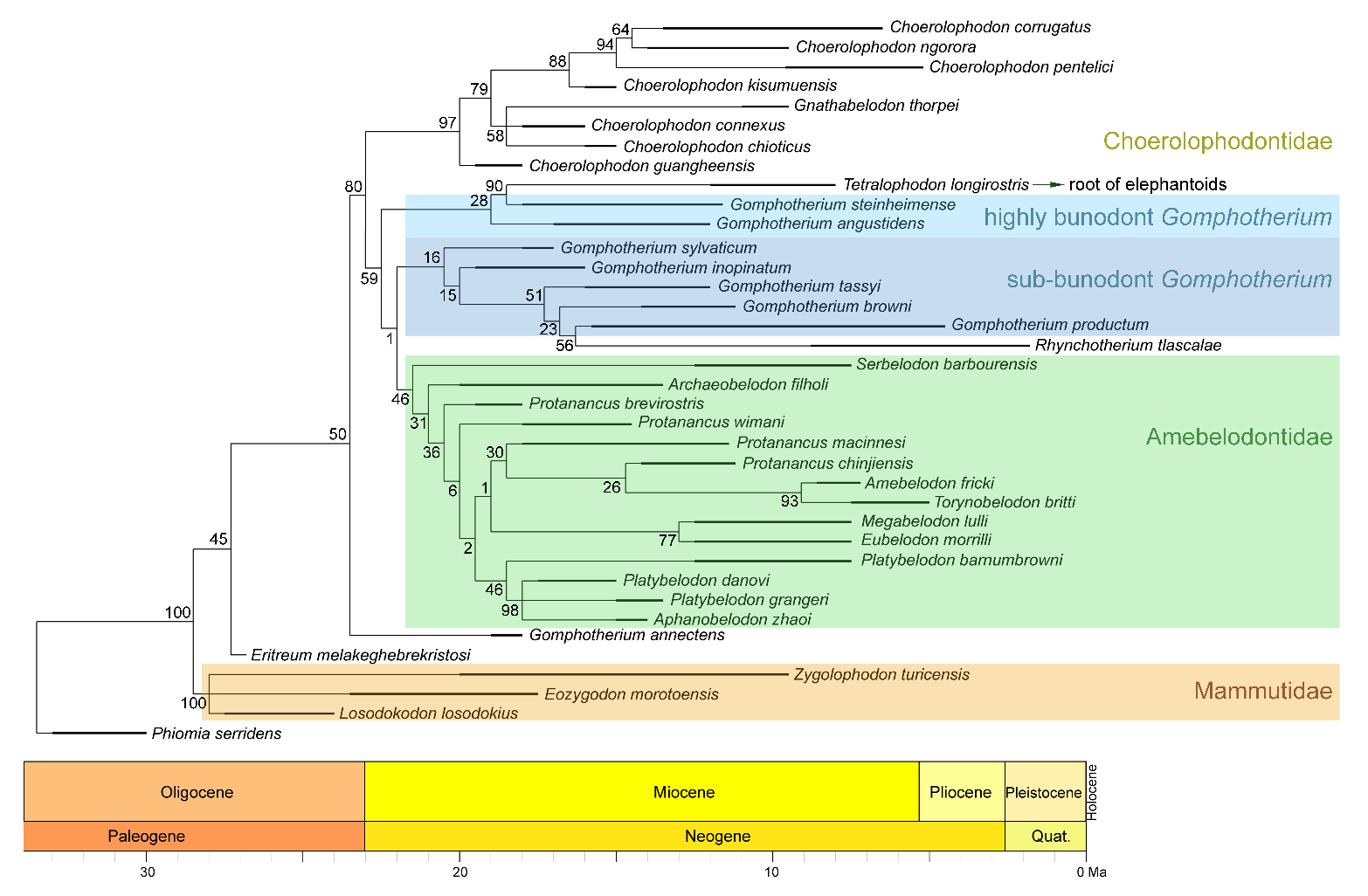

Figure S1 Phylogenetic reconstruction of longirostrine bunodont elephantiforms and mammutids using the maximum parsimony method (tree length, 253; CI, 0.470; RI, 0.779). The node support (the number at each node) was calculated by symmetric resampling with 0.33 change probability (1000 replicates).

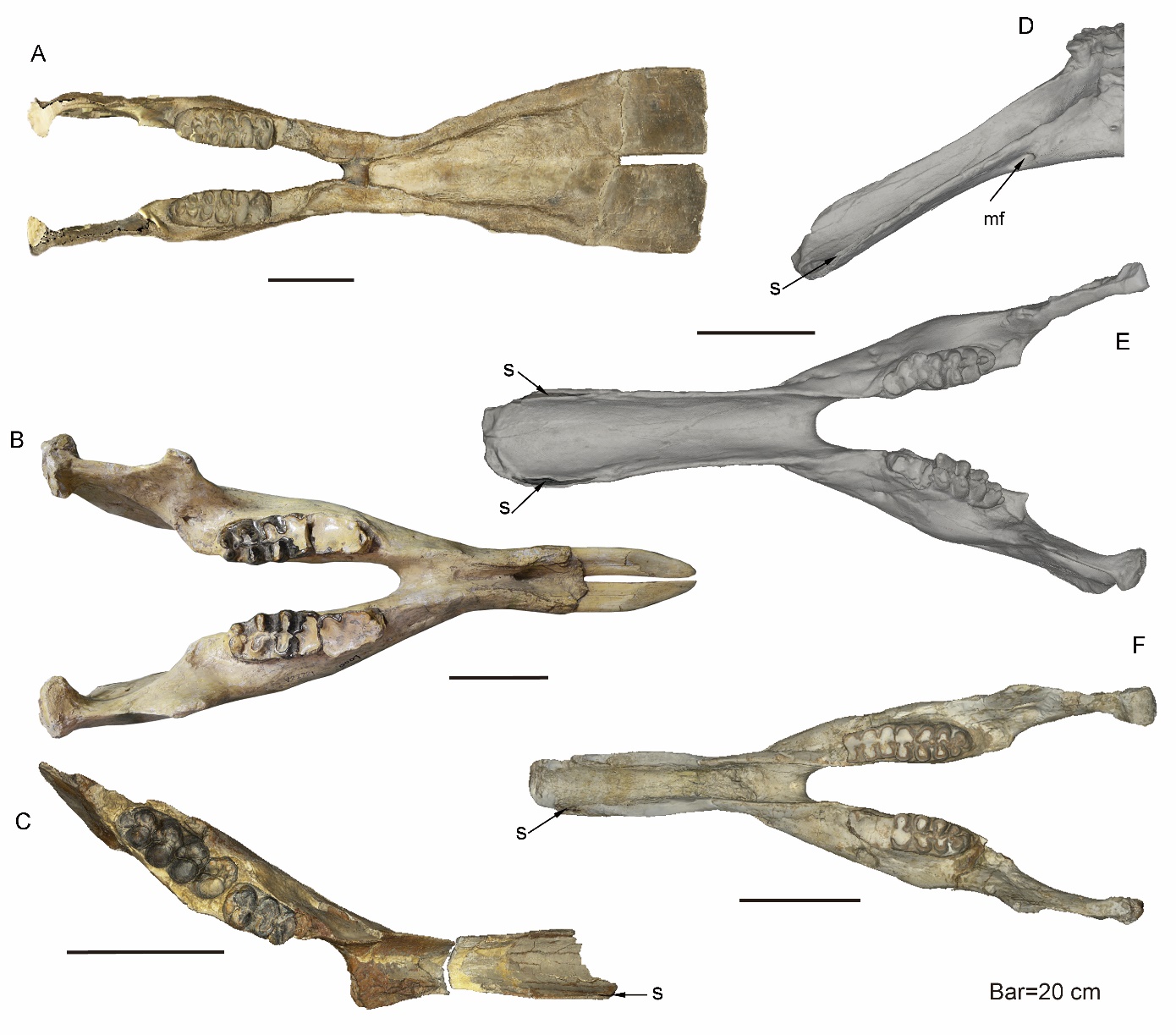

Figure S2. Mandible of longirostrine bunodont elephantiforms. (**A**) *Platybelodon grangeri*, HMV 0939, from Zengjia Fauna, Linxia Basin. (**B**) *Gomphotherium tassyi*, IVPP V22781, from Hejiagou Fauna, Tongxin region. (**C**) *Choerolophodon connexus*, IVPP V8567, from Halamagai Fauna, Junggar Basin. (**D–F**) *Choerolophodon chioticus*, from Lower and Middle Miaoerling Faunas, Tongxin region, including IVPP V23457, male, lateral view [D], showing the large, tube-like anterior mental foramen, and dorsal view [E] and IVPP V23213 [F], female, dorsal view. Anatomic abbreviations: mf, mental foramen; s, slit for holding keratinous cutting plates in *Choerolophodon*.

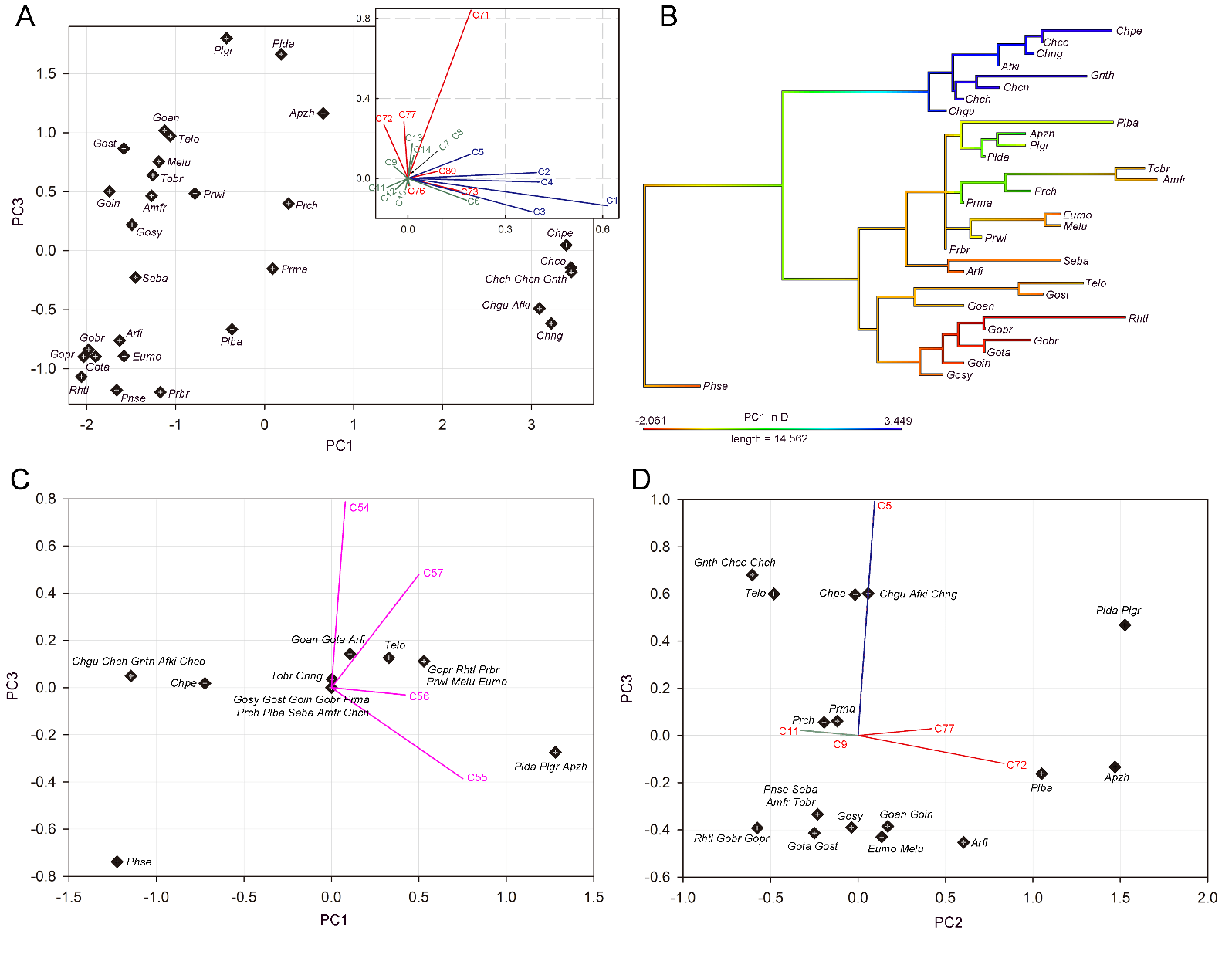

Figure S3. Character-combines in relation to feeding behavior in trilophodont longirostrine bunodont elephantiforms. (**A**) PC1–PC3 plan of character-combine of tusks and mandibular symphysis; (**B**) PC1 scores of A that are mapped on the phylogenetic tree (mammutids and stem elephantimorphs excluded); (**C**) PC1–PC3 plan of character-combine of narial region; (**D**) PC2–PC3 plan of character combine in relation to horizontal cutting behavior. In A, C, and D, character loads were shown as line segments. Notes: character load vectors were marked in purple, narial region; red, mandible; blue, upper tusk; and green, mandibular tusk, colors, respectively. Taxa abbreviation: Afki, *Afrochoerodon kisumuensis*; Amfr, *Amebelodon fricki*; Apzh, *Aphanobelodon zhaoi*; Arfi, *Archoerobelodon filholi*; Chch, *Choerolophodon chioticus*; Chco, *Choerolophodon corrugatus*; Chcn, *Choerolophodon connexus*; Chgu, *Choerolophodon guangheensis*; Chng, *Choerolophodon ngorora*; Eumo, *Eubelodon morrilli*; Gnth, *Gnathabelodon thorpei*; Goan, *Gomphotherium angustidens*; Gobr, *Gomphotherium browni*; Goin, *Gomphotherium inopinatum*; Gost, *Gomphotherium steinheimense*; Gopr, *Gomphotherium productum*; Gosy, *Gomphotherium sylvaticum*; Gota, *Gomphotherium tassyi*; Melu, *Megabelodon lulli*; Phse, *Phiomia serridens*; Plba, *Platybelodon barnumbrowni*; Plda, *Platybelodon danovi*; Plgr, *Platybelodon grangeri*; Plte, *Platybelodon tetralophus*; Plto, *Platybelodon tongxinensis*; Prch, *Protanancus chinjiensis*; Prbr, *Protanancus brevirostris*; Prma, *Protanancus* *macinnesi*; Prwi, *Protanancus wimani*; Rhtl, *Rhynchotherium tlascalae*; Seba, *Serbelodon barbourensis*; Telo, *Tetralophodon longirostris*; *Tobr, Torynobelodon britti*.

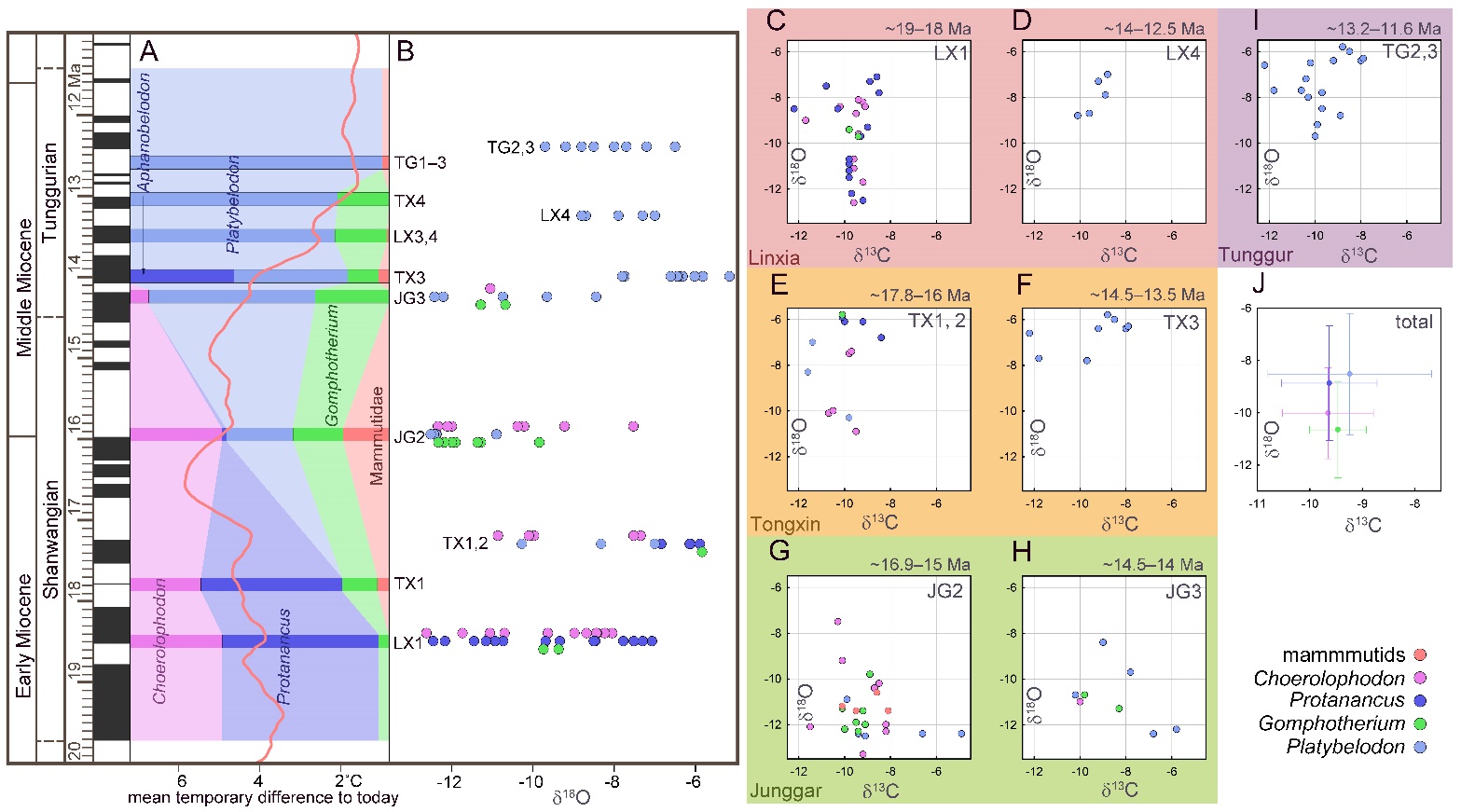

Figure S4. Relative abundance and tooth enamel δ^18^O values of fossil elephantiforms from the Shanwangian and Tunggurian stages (~19–11.5 Ma) in northern China. (**A**) Relative abundances of the four elephantiform families, including Choerolophodontidae, only represented by *Choerolophodon* (pink); Amebelodontidae, represented by *Protanancus*, *Aphanobelodon* (dark blue), and *Platybelodon* (light blue); “Gomphotheriidae”, only represented by *Gomphotherium* (green); and Mammutidae (red). Horizontal bars represent the average ages of the fossil assemblages. The ages were determined by palaeomagnetism (Table S1). The red curve shows the global benthic foraminifer oxygen isotope curve, which represents the global temperature (after Westerhold^51^). (**B**) Tooth enamel stable oxygen isotopic compositions of various gomphothere taxa. Each circle represents a single data point. (**C–I**) Plots of gomphothere tooth enamel stable carbon and oxygen isotopes in different fossil assemblages, including Dalanggou [C] and Laogou [D], Linxia Basin; lower Miaoerling [E] and upper Miaoerling [F], Tongxin Region; Halamagai [G] and Kekemaideng [H], Junggar Basin; and Moergen + Tamuqin [I], Tunggur region. (**J**) Isotopic data statistics from all gomphothere samples. The circles represent the average values, and the error bars represent the standard deviations. Abbreviations for fossil assemblages: JG1, Top Suosuoquan Fauna; JG2, Halamagai Fauna; JG3, Kekemaideng Fauna; JG4, Dingshanyanchi I Fauna; LX1, Dalanggou Fauna; LX2, Shinanu Fauna; LX3, Zengjia Fauna; LX4, Laogou Fauna; TG1, Tairum Fauna; TG2, Moergen Fauna; TG3, Tamuqin Fauna; TX1, Lower Miaoerling (Miaoerling A/B) Fauna; TX2, Middle Miaoerling (Miaoerling C) Fauna; TX3, Upper Miaoerling (Miaoerling D/E) Fauna; TX4, Heijiagou Fauna.

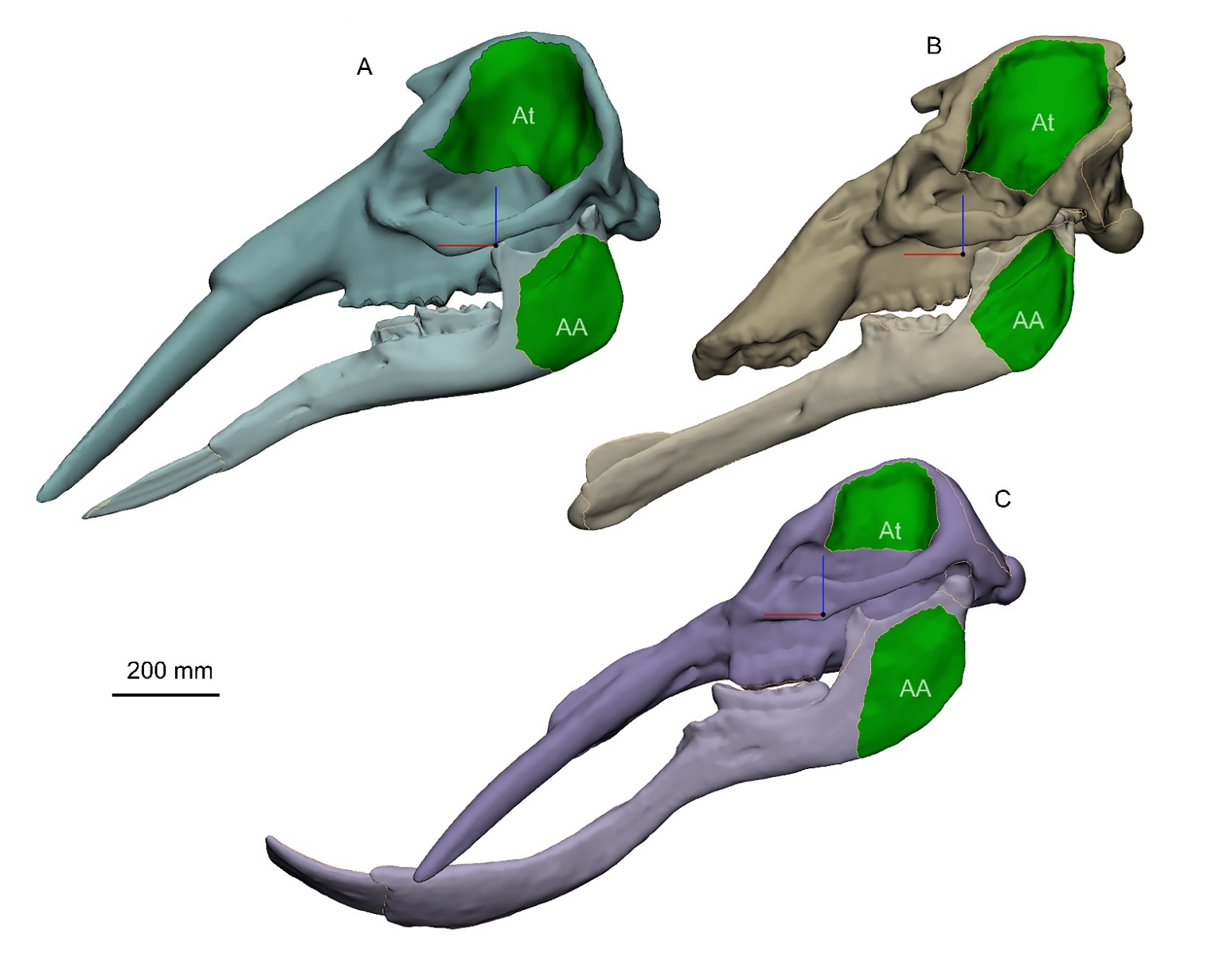

Figure S5. Geometric skull models for FE analysis of three representative bunodont elephantiforms, including *Gomphotherium* (**A**), *Choerolophodon* (**B**), and *Platybelodon* (**C**), generated by Materialise 3-matic Research (V12.0). For comparative purposes, the three bony parts of mandibles (excluding the tusk or cutting plate) were scaled to the same volume as those of *Choerolophodon*, and the other parts (cranium and tusk or cutting plate) were also scaled to the bony mandible of each model. The green areas indicate the surfaces for attaching *temporalis* (*At*), and *superficial* *masseter* + *zygomaticomandibularis* + *pterygoideus internus* (*AA*), of which the areas were calculated for muscle force estimation. Note that the muscle forces of *Choerolophodon* were used in FE analyses, and those of *Gomphotherium* and *Platybelodon* were assigned the same magnitude as the muscle forces of *Choerolophodon* for comparative purposes.

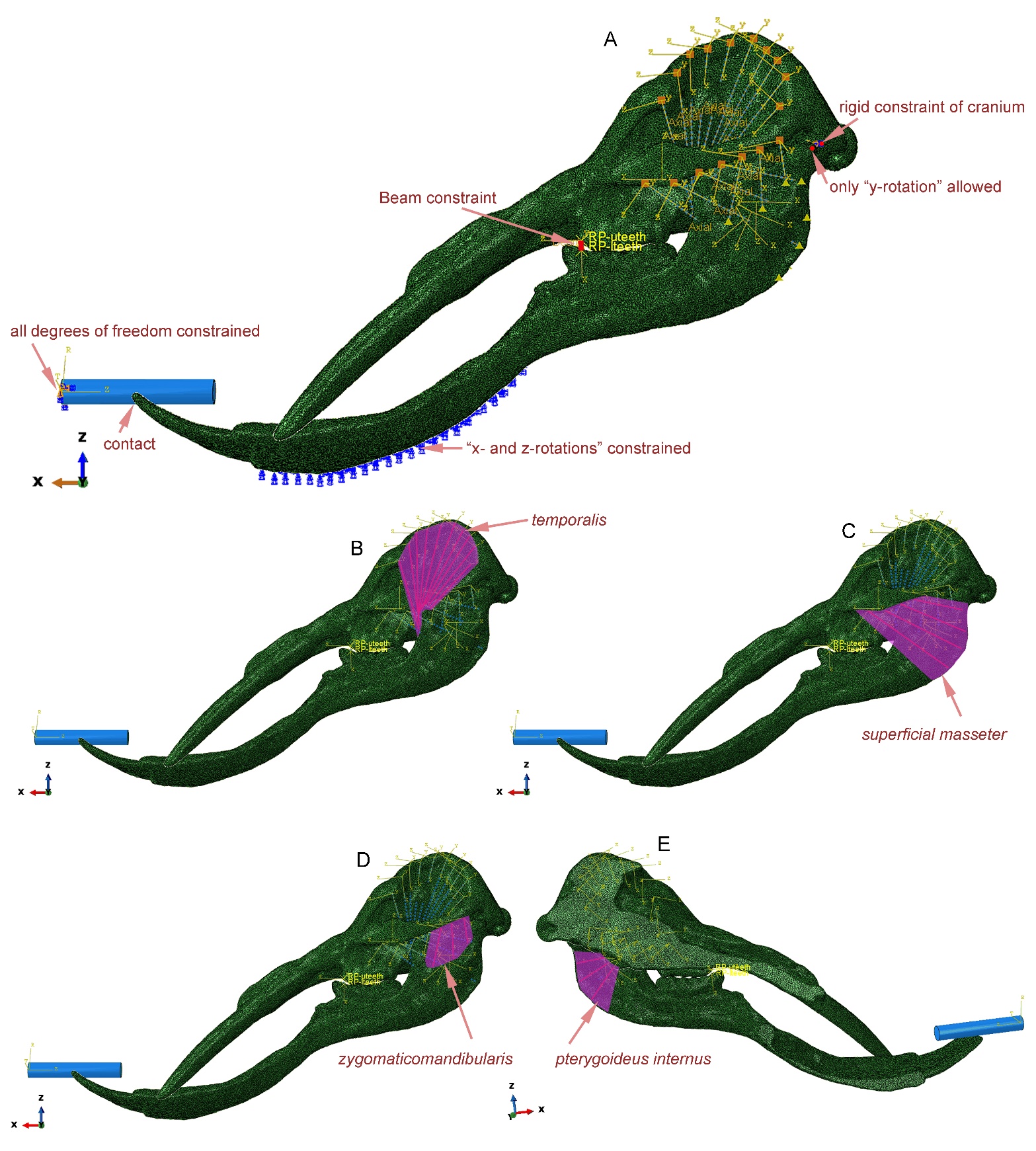

Figure S6. Mechanical settings of *Platybelodon* horizontal twig-cutting modelling; the blue cylinder represents the twig model. (**A**) Constraints and boundary conditions of the model. (**B**) Modelling of the *temporalis* muscle; the muscle forces were exerted on the red axes. (**C**) Modelling of the *superficial* *masseter* muscle. (**D**) Modelling of the *zygomaticomandibularis* muscle. (**E**) Modelling of the *pterygoideus internus* muscle.

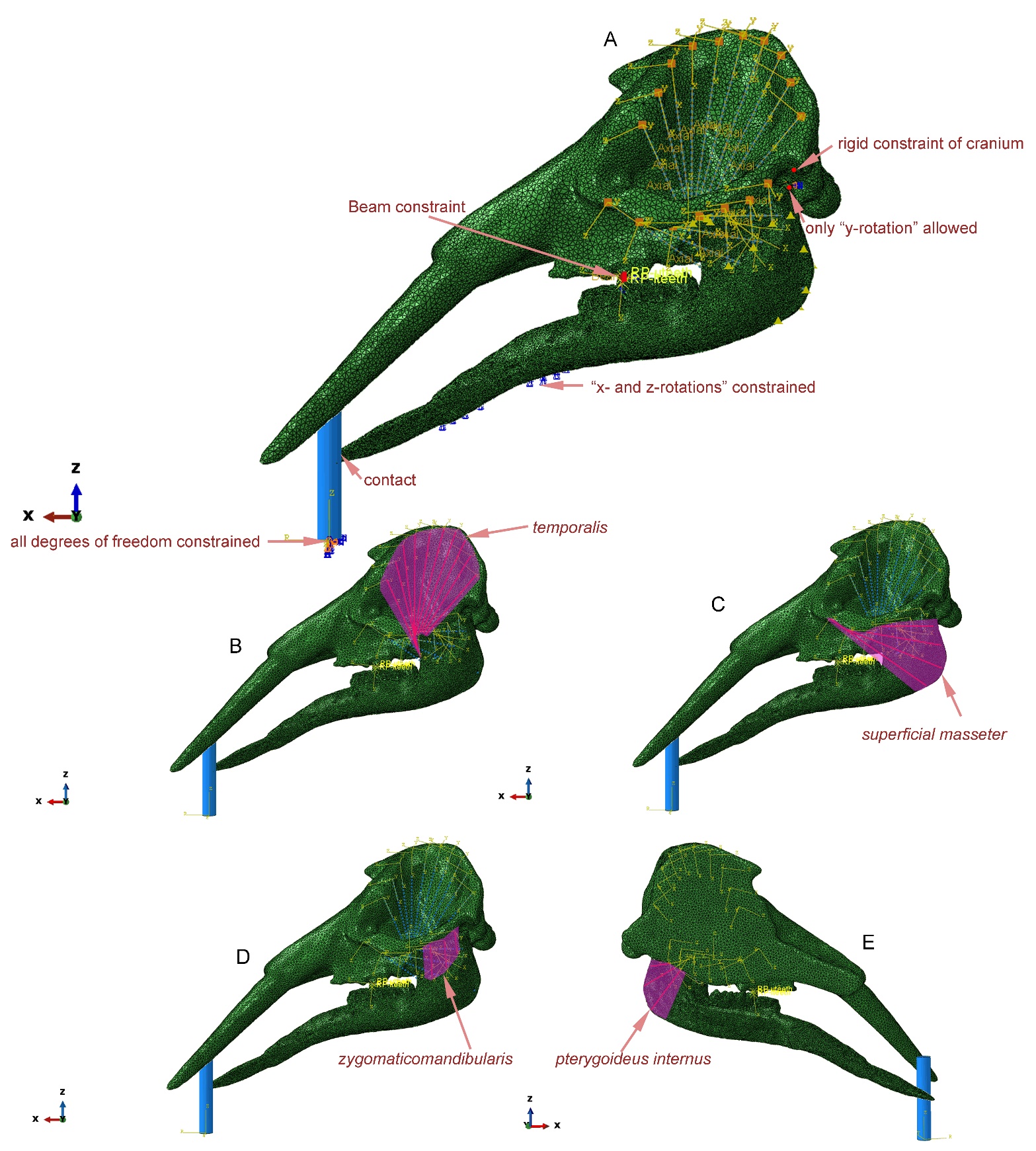

Figure S7. Mechanical settings of *Gomphotherium* vertical twig-cutting modelling. (**A**) Constraints and boundary conditions of the model. (**B**) Modelling of the *temporalis* muscle. (**C**) Modelling of the *superficial* *masseter* muscle. (**D**) Modelling of the *zygomaticomandibularis* muscle. (**E**) Modelling of the *pterygoideus internus* muscle.

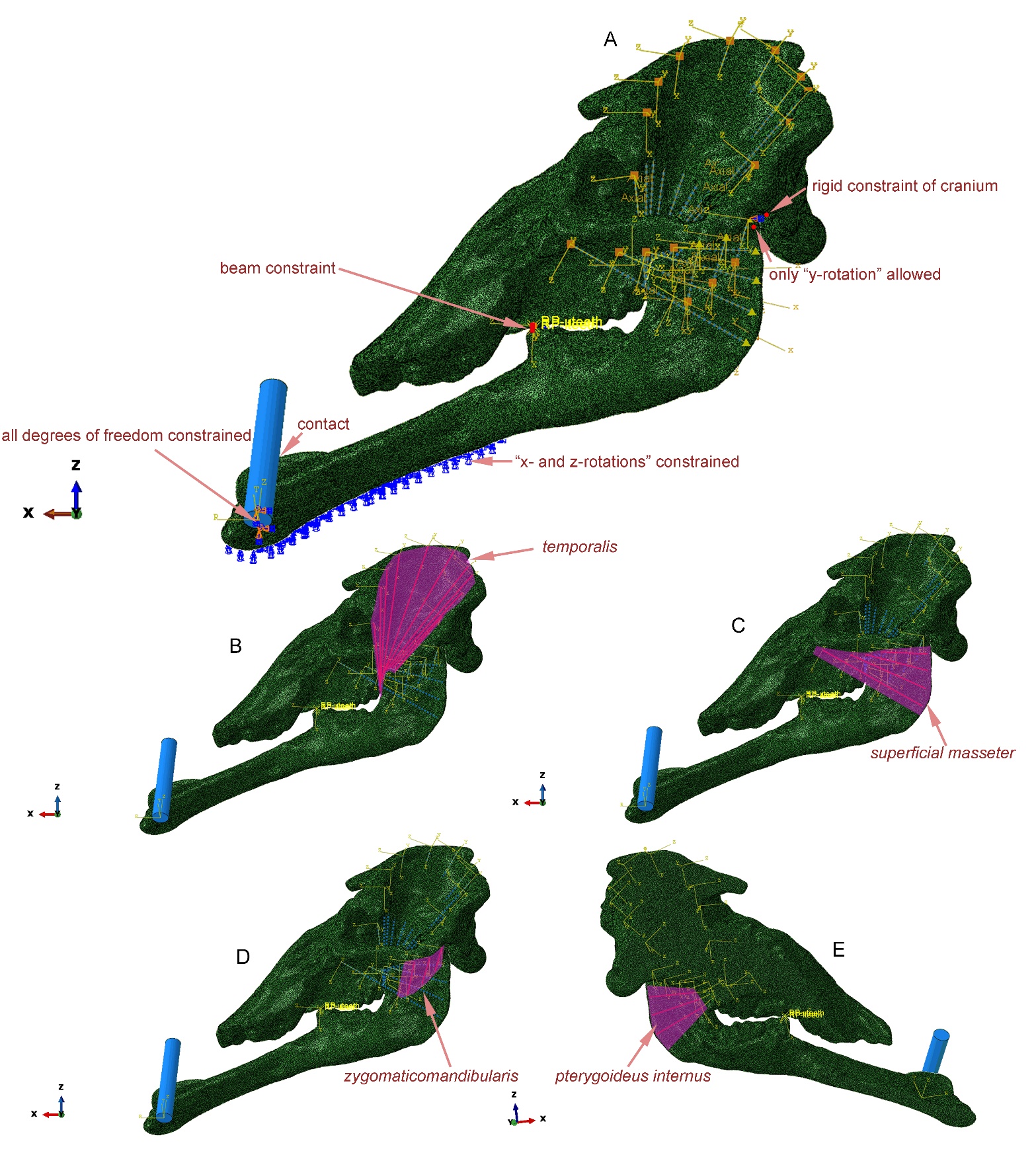

Figure S8. Mechanical settings of *Choerolophodon* oblique twig-cutting modelling. (**A**) Constraints and boundary conditions of the model. (**B**) Modeling of the *temporalis* muscle. (**C**) Modelling of the *superficial* *masseter* muscle. (**D**) Modelling of the *zygomaticomandibularis* muscle. (**E**) Modelling of the *pterygoideus internus* muscle.

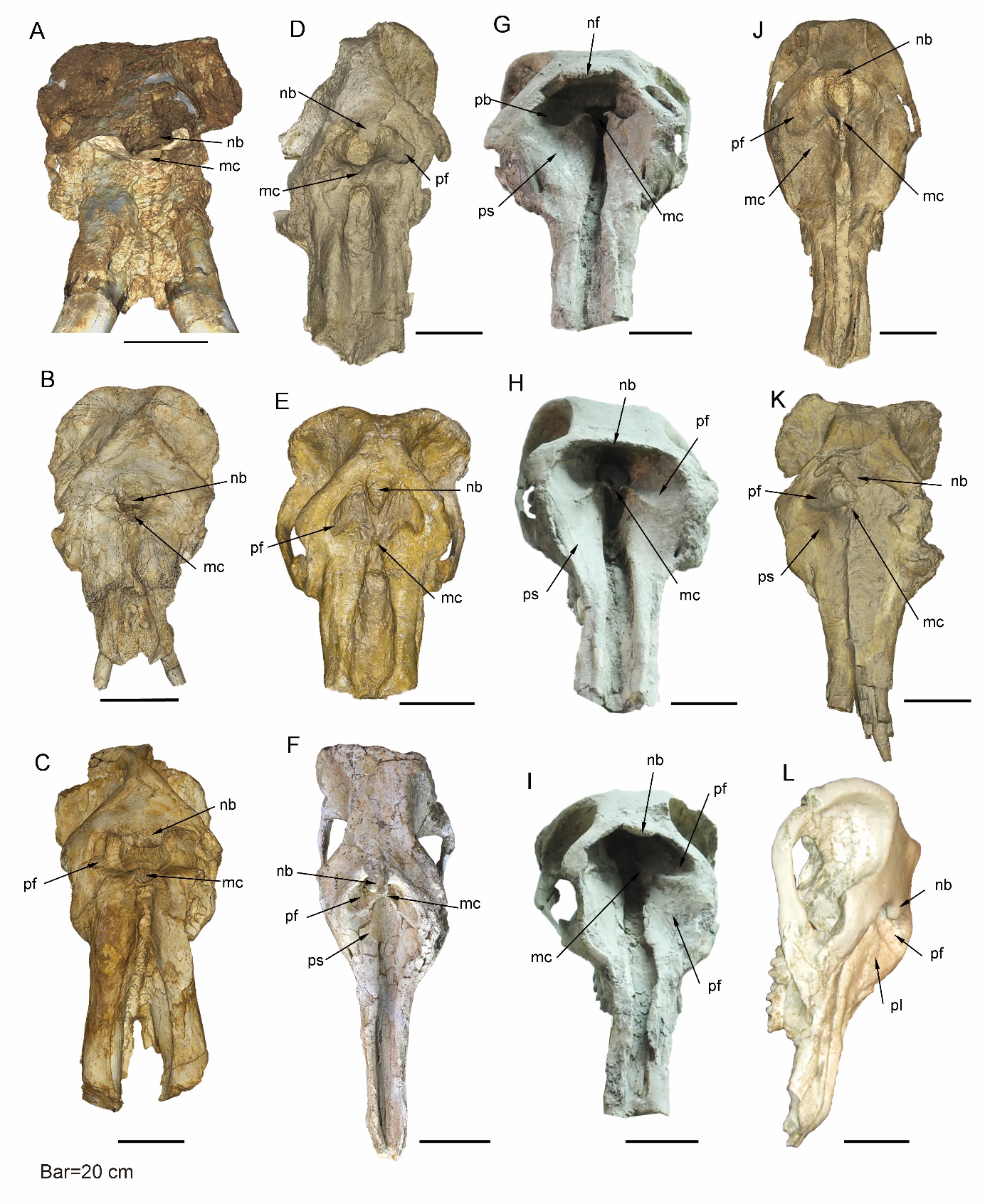

Figure S9. Narial region of longirostrine bunodont elephantiforms. (**A–B**), *Choerolophodon guangheensis* showing the large nasal bone processes and insertion for mesethmoid cartilage, lacking perinasal fossae; IVPP V17685 [A], type specimen, male, from Dalanggou Fauna, Linxia Basin; IVPP V31268 [B], female, from Lower Miaoerling Fauna, Tongxin region. (**C**) *Choerolophodon chioticus*, IVPP V23457, male, from Middle Miaoerling Fauna, Tongxin region, showing a large nasal bone process and mesethmoid cartilage insertion, with very small perinasal fossae. (**D–E**) *Gomphotherium tassyi* showing a moderately developed nasal bone processes and mesethmoid cartilage insertion, and well-developed perinasal fossae; HMV 0028 [D], type specimen, from Zengjia fauna, Linxia; IVPP V22780 [E], from Hejiagou fauna, Tongxin region. (**F**) *Aphanobelodon zhaoi*, HMV 1880, type specimen, from Upper Miaoerling fauna, Tongxin, showing a small nasal bone process and mesethmoid cartilage insertion, well-developed perinasal fossae, and initially developed prenasal slope. (**G–J**) *Platybelodon grangeri*, showing the very small nasal bone process and mesethmoid cartilage insertion, well-developed perinasal fossae and prenasal slope, including HMV 1841 [G], HMV 1840 [H], HMV 0940 [I], and HMV 0939 [J], from Zengjia Fauna, Linxia Basin. (**K**) *Platybelodon tongxinensis*, from Lintong region showing the small nasal bone process and mesethmoid cartilage insertion, well-developed perinasal fossae, and initially developed prenasal slope. (**L**) *Platybelodon tetralophus*, AMNH 26460, from Tamuqin Fauna, Tunggur region, showing a small nasal bone process, and well-developed perinasal fossae and prenasal slope. Anatomic abbreviations: ce, cutting edge of the distal mandibular tusk in *Platybelodon*; nb, nasal process of nasal bone; mc, slit or groove for mesethmoid cartilage insertion (white in colour); pf, perinasal fossa; ps, prenasal slope.

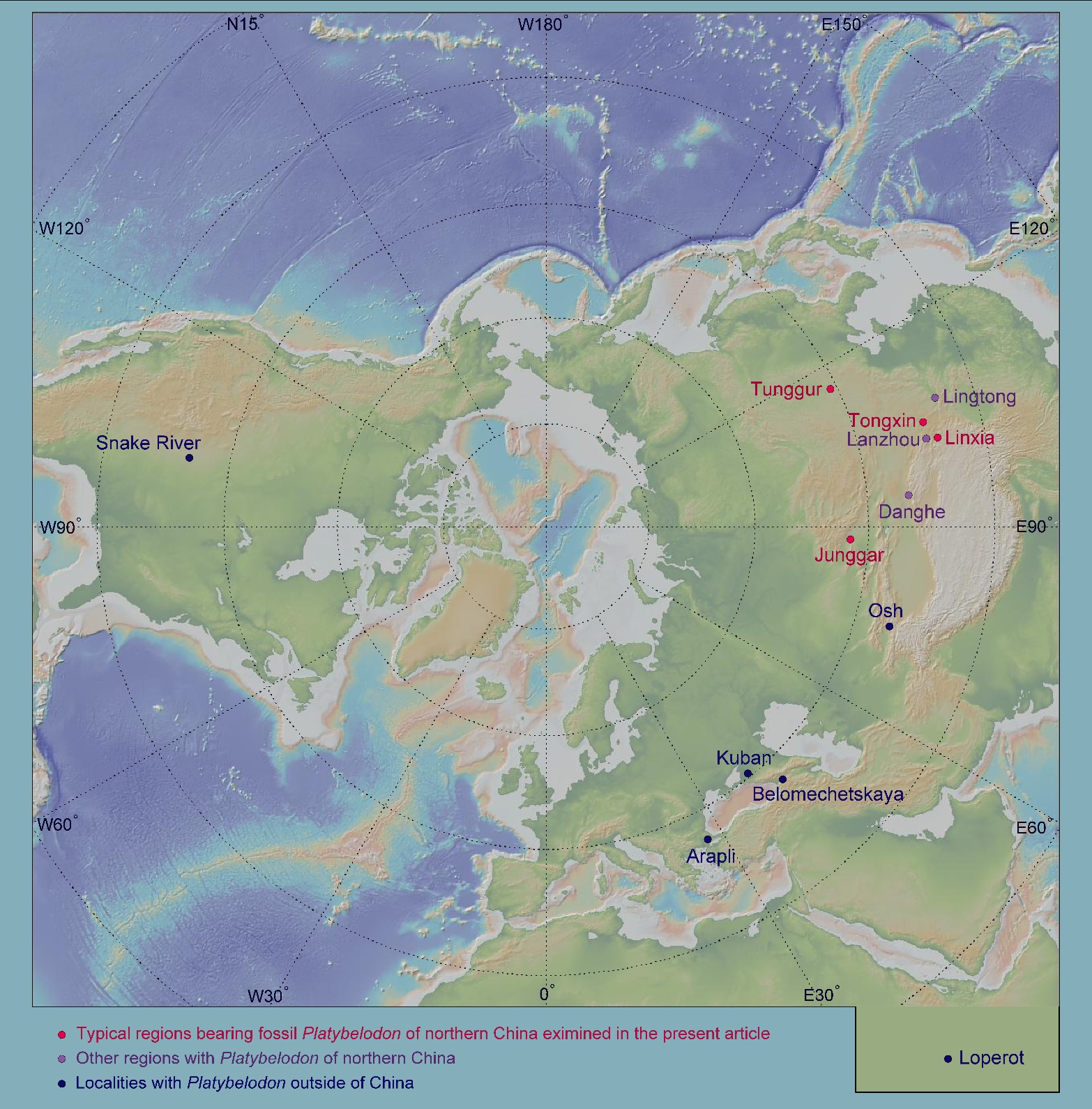

Figure S10. Geographic distribution of *Platybelodon* worldwide. The red circles represent the four regions examined in the present article, the light purple circles show the other regions with *Platybelodon* in northern China, and the dark blue circles denote the *Platybelodon* localities outside of China. This geographic map was generated by GeoMapApp (V3.6.15) (https://www.geomapapp.org/).

#### Supplementary Tables

Table S1. Age estimation of different faunas of the four regions, dating after Sun^6^, Wang^7^, Wang et al.^4^ and Qiu et al.^5^.

| region | locality | beginning age (Ma) | ending age (Ma) | average age (Ma) | references |
| --- | --- | --- | --- | --- | --- |
| Linxia Basin | Dalanggou (LX1) | 19 | 18 | 18.5 | Sun 2014, reinterpreted |
| Linxia Basin | Shinanu (LX2) | 17 | 15.2 | 16.1 | Sun 2014, reinterpreted |
| Linxia Basin | Zengjia (LX3) | 14.5 | 14 | 14.25 | Sun 2014, reinterpreted |
| Linxia Basin | Laogou (LX4) | 14 | 12.5 | 13.25 | Sun 2014, reinterpreted |
| Tongxin Region | Lower Miaoerling (TX1) | 17.8 | 17 | 17.4 | Wang 2021, reinterpreted |
| Tongxin Region | Middle Miaoerling (TX2) | 16.6 | 15 | 15.8 | Wang 2021, reinterpreted |
| Tongxin Region | Upper Miaoerling (TX3) | 14.5 | 13.5 | 14 | Wang 2021, reinterpreted |
| Tongxin Region | Heijiagou (TX4) | 13.4 | 12.7 | 13.05 | estimated, based on Wang 2021* |
| Junggar Basin | top Suosuoquan (JG1) | 18.8 | 18.4 | 18.6 | Wang et al., 2022 |
| Junggar Basin | Halamagai (JG2) | 16.9 | 15 | 15.95 | Wang et al., 2022 |
| Junggar Basin | Kekemaideng (JG3) | 14.5 | 14 | 14.25 | Wang et al., 2022 |
| Junggar Basin | Dingshanyanchi I (JG4) | 13 | 12.6 | 12.8 | Wang et al., 2022 |
| Tunggur region | Tairum (TG1) | 13.6 | 13.2 | 13.4 | Qiu et al., 2013 |
| Tunggur region | Moergen (TG2) | 13.2 | 12 | 12.6 | Qiu et al., 2013 |
| Tunggur region | Tamuqin (TG3) | 12 | 11.6 | 11.8 | Qiu et al., 2013 |

*Hejiagou is located at Zhongning county, and the strata are correlated to the upper part of the Miaoerling Section, both belonging to the Zhangenbao Formation.

Table S2 Geometric and mesh parameters of the FE models.

| Models | element type | element number | volume (mm^3^) | scale factor (%) |
| --- | --- | --- | --- | --- |
| *Choerolophodon*, cranium | C3D4 | 306068 | 27763213.46 | 1 |
| *Choerolophodon*, mandible | C3D4 | 128982 | 6563708.01 | 1 |
| *Choerolophodon*, keratinous sheet | C3D4 | 27648 | 80054.10 | 1 |
| *Platybelodon*, cranium | C3D4 | 96734 | 15567961.81 | 84.23 |
| *Platybelodon*, mandible | C3D4 | 110010 | 6563703.39 | 84.23 |
| *Platybelodon*, mandibular tusk | C3D4 | 30416 | 1255711.19 | 84.23 |
| *Gomphotherium*, cranium | C3D4 | 42666 | 28441280.58 | 99.11 |
| *Gomphotherium*, mandible | C3D4 | 92780 | 6563709.82 | 99.11 |
| *Gomphotherium*, mandibular tusk | C3D4 | 18564 | 438420.77 | 99.11 |
| twig | C3D8R | 106650 | 589048.62 | ― |

Table S3 Material properties in the FE models, after Drake et al.^28^, Huo et al.^29^, and Risbrudt et al.^30^

|  | Elasticity parameters |  |  |
| --- | --- | --- | --- |
| Materials | Young's modulus (MPa) | Poisson's ratio |  |
| bone | 15000 | 0.28 |  |
| dentine (tusk) | 29500 | 0.44 |  |
| keratinous integument (keratin) | 2000 | 0.3 |  |
| wood (wet red pine) subscript: L: axial; R: radial; T, tangential | orthotropic elastoplasticity parameters |  |  |
| direct modulus (MPa) | E_L_ | E_R_ | E_T_ |
|  | 9680 | 425.92 | 851.84 |
| shearing modulus (MPa) | G_LR_ | G_LT_ | G_RT_ |
|  | 929.28 | 784.08 | 106.48 |
| Poisson's ratio | μ_LR_ | μ_LT_ | μ_RT_ |
|  | 0.347 | 0.315 | 0.408 |
| yield stress (MPa) | f_0_ |  |  |
|  | 18.8 |  |  |
| yield ratio | R_LL_ | R_RR_ | R_TT_ |
|  | 1 | 0.0957 | 0.0957 |
|  | R_LT_ | R_RT_ | R_LR_ |
|  | 0.4422 | 0.4422 | 0.4422 |

Table S4 Muscle forces estimation of FE models

|  | At, temporal fossa area (mm^2^) | AA, *masseter*-complex attaching area (mm^2^) | force on each axial connector of the *temporalis* (N) | force on each axial connector of the *masseter*-complex (N) | external force (N) |
| --- | --- | --- | --- | --- | --- |
| *Choerolophodon* | 103332.7665 | 49608.1101 | 3099.982995 | 1488.243303 | 5000 |
| *Platybelodon** | 35435.547 | 51968.0569 | 3099.982995 | 1488.243303 | 5000 |
| *Gomphotherium** | 89298.493 | 56203.2467 | 3099.982995 | 1488.243303 | 5000 |

*Scaled to the same volume as *Choerolophodon*, and the uniform muscle forces estimated from *Choerolophodon* model were used, see the Methods part

#### List of the supplementary data, codes, videos, and 3D digital models

##### Supplementary data

Data S1 Morphological characters and data set for phylogenetic analyses, characters see Appendix S1.

Data S2 Elephantiform specimens counted in the present article, are housed in IVPP, HPM, and AMNH.

Data S3 Original measurements of elephantiform tooth enamel isotope ratios analyses in the present article.

##### Supplementary codes

Code S1 Script file for Bayesian total-evidence dating analysis (.nex file).

Code S2 Script auxiliary file for most parsimonious analysis, applying for some ordered and irreversible characters (characters 19–24 in the TNT program, please note that these character numbers were automatically assigned by TNT file, and begins from character “0” rather than “1”, therefore, these characters in supplementary Appendix and Data S1 were characters 20–25) (.tnt file).

Code S3 Script file for most parsimonious analysis (.tnt file).

Code S4 FEA setting file, for distal force tests (this file can be directly submitted in the Abaqus 6.14).

Code S5 FEA setting file, for distal twig cutting tests (this file can be directly submitted in the Abaqus 6.14).

##### Supplementary Movies

Movie S1 FE modeling of *Platybelodon* distal force test, color map showing the von Mises stress.

Movie S2	FE modeling of *Gomphotherium* distal force test, color map showing the von Mises stress.

Movie S3	FE modeling of *Choerolophodon* distal force test, color map showing the von Mises stress.

Movie S4 FE modeling of *Platybelodon* vertical twig cutting test, the total model, color map showing the von Mises stress.

Movie S5 FE modeling of *Platybelodon* vertical twig cutting test, the twig model (facing to the surface that is in contact with the tusk), color map showing the equivalent plastic strain.

Movie S6 FE modeling of *Platybelodon* oblique twig cutting test, the total model, color map showing the von Mises stress.

Movie S7 FE modeling of *Platybelodon* oblique twig cutting test, the twig model (facing to the surface that is in contact with the tusk), color map showing the equivalent plastic strain.

Movie S8 FE modeling of *Platybelodon* horizontal twig cutting test, the total model, color map showing the von Mises stress.

Movie S9 FE modeling of *Platybelodon* horizontal twig cutting test, the twig model (facing to the surface that is in contact with the tusk), color map showing the equivalent plastic strain.

Movie S10 FE modeling of *Gomphotherium* vertical twig cutting test, the total model, color map showing the von Mises stress.

Movie S11 FE modeling of *Gomphotherium* vertical twig cutting test, the twig model (facing to the surface that is in contact with the tusk), color map showing the equivalent plastic strain.

Movie S12 FE modeling of *Gomphotherium* oblique twig cutting test, the total model, color map showing the von Mises stress.

Movie S13 FE modeling of *Gomphotherium* oblique twig cutting test, the twig model (facing to the surface that is in contact with the tusk), color map showing the equivalent plastic strain.

Movie S14 FE modeling of *Gomphotherium* horizontal twig cutting test, the total model, color map showing the von Mises stress.

Movie S15 FE modeling of *Gomphotherium* horizontal twig cutting test, the twig model (facing to the surface that is in contact with the tusk), color map showing the equivalent plastic strain.

Movie S16 FE modeling of *Choerolophodon* oblique twig cutting test, the total model, color map showing the von Mises stress.

Movie S17 FE modeling of *Choerolophodon* oblique twig cutting test, the twig model (facing to the surface that is in contact with the cutting plate), color map showing the equivalent plastic strain.

Movie S18 FE modeling of *Choerolophodon* horizontal twig cutting test, the total model, color map showing the von Mises stress.

Movie S19 FE modeling of *Choerolophodon* horizontal twig cutting test, the twig model (facing to the surface that is in contact with the cutting plate), color map showing the equivalent plastic strain.

##### Supplementary 3D models

3D model S1 *Platybelodon* cranium model (.stl file)

3D model S2 *Platybelodon* mandible model (.stl file)

3D model S3 *Gomphotherium* cranium model (.stl file)

3D model S4 *Gomphotherium* mandible model (.stl file)

3D model S5 *Choerolophodon* cranium model (.stl file)

3D model S6 *Choerolophodon* mandible model (.stl file)
